# Aging and memory are altered by genetically manipulating lactate dehydrogenase in the neurons or glia of flies

**DOI:** 10.1101/2022.10.19.512981

**Authors:** Ariel K. Frame, J. Wesley Robinson, Nader H. Mahmoudzadeh, Jason M. Tennessen, Anne F. Simon, Robert C. Cumming

## Abstract

The astrocyte-neuron lactate shuttle hypothesis posits that glial-generated lactate is transported to neurons to fuel metabolic processes required for long-term memory. Although studies in vertebrates have revealed that lactate shuttling is important for cognitive function, it is uncertain if this form of metabolic coupling is conserved in invertebrates or is influenced by age. Lactate dehydrogenase (Ldh) is a rate limiting enzyme that interconverts lactate and pyruvate. Here we genetically manipulated expression of *Drosophila melanogaster* lactate dehydrogenase (dLdh) in neurons or glia to assess the impact of altered lactate metabolism on invertebrate aging and long-term courtship memory at different ages. We also assessed survival, negative geotaxis, brain neutral lipids (the core component of lipid droplets) and brain metabolites. Both upregulation and downregulation of dLdh in neurons resulted in decreased survival and memory impairment with age. Glial downregulation of dLdh expression caused age-related memory impairment without altering survival, while upregulated glial dLdh expression lowered survival without disrupting memory. Both neuronal and glial dLdh upregulation increased neutral lipid accumulation. We provide evidence that altered lactate metabolism with age affects the tricarboxylic acid (TCA) cycle, 2-hydroxyglutarate (2HG), and neutral lipid accumulation. Collectively, our findings indicate that the direct alteration of lactate metabolism in either glia or neurons affects memory and survival but only in an age-dependent manner.

**Author Summary:** The brain is composed of metabolically demanding cell types that must remain functional throughout life to maintain cognitive tasks like memory formation. How brain metabolism varies between cell-types across the lifespan is uncertain. It has been suggested that in vertebrate brains glia breakdown sugars to produce lactate and shuttle it to neurons to support long-term memory and survival. Yet, this phenomenon has not been directly tested in invertebrates. In *Drosophila melanogaster* flies, we genetically manipulated lactate dehydrogenase, a central enzyme in lactate metabolism, specifically within the adult brain. We discovered that shifting lactate metabolism to either higher or lower levels in neurons caused decreased survival and worse long-term memory with age. In contrast, we found increased glial lactate metabolism worsened survival, whereas decreased glial lactate metabolism impaired long-term memory in aged flies. Both increased glial or neuronal lactate metabolism led to increased accumulation of metabolites, such as 2-hydroxyglutarate, mitochondrial metabolites, and neutral lipids, in aged fly brains. These findings provide evidence for the first time in invertebrates that lactate and memory are connected and altered lactate metabolism in neurons and glia differentially impact survival and memory, but only in an age-dependent manner.

**Graphical Abstract:** 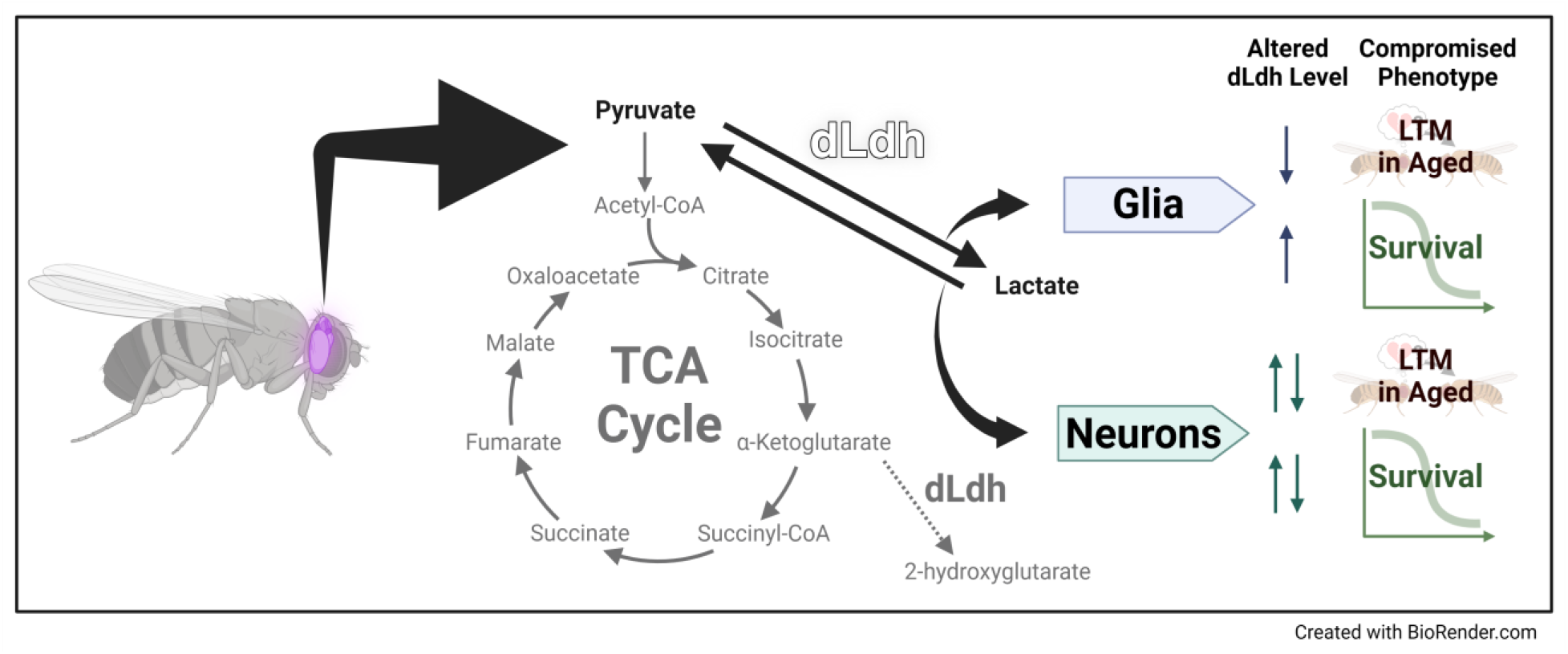

## Introduction

The brain is an energetically demanding organ relative to other tissues, a phenomenon found in multiple species from humans to flies [1–4]. Various metabolites can be utilized to meet the energy requirements of neural cells, although the preferred metabolite may differ by cell-type, brain region, age, and behavioural context. Over the last two decades there has been growing recognition that lactate, the end product of glycolysis, serves many functions, including acting as a source of energy, a signaling molecule, and even as an epigenetic regulator [5].

Multiple studies on the brains of vertebrates have lent support to the hypothesis that astrocytes supply neurons with lactate for their energetic needs [6,7,16,17,8–15]. This type of metabolic coupling is commonly referred to as the astrocyte-neuron lactate shuttle (ANLS) [18,19]. Despite the accumulation of a large body of evidence supporting the existence of an ANLS, there is still controversy surrounding the physiological role and importance of the ANLS [20–26]. How the ANLS contributes to behaviour has yet to be fully elucidated. Most studies investigating the role that the ANLS plays in behaviour have focused on mammals. However, astrocyte function is also conserved in invertebrates [27–30]. In fact, evidence for a glia-neuron metabolic coupling in invertebrates actually preceded the discovery of the ANLS [31–33]. Recent studies suggest that invertebrate animals such as *Drosophila melanogaster* rely on glia to produce and release lactate, through monocarboxylate transporters, to support neuronal survival, synaptic function, axonal regeneration, and synthesis of fatty acids in a manner analogous to the ANLS in mammals [34–38]. Therefore, *D. melanogaster* serves as a good model for understanding the role of glia-neuron lactate shuttling in central nervous system (CNS) function.

The ANLS has been implicated in multiple cognitive processes in mammals [39–47]. In contrast, relatively few studies have examined the effect of glia-neuron metabolic coupling on cognition in *D. melanogaster* [48–50]. Two recent findings suggest that *D. melanogaster,* in certain contexts, require glia to provide glucose or ketone bodies to fuel neurons, irrespective of lactate, and facilitate memory formation [48,49]. Moreover, findings by our laboratory and others have provided evidence that glia-neuron lactate shuttling in *D. melanogaster* impacts aging and neurodegeneration [36,51,52]. In addition, glia have been shown to contribute to age-related memory impairment [53,54]. Nonetheless, the contribution of glia-neuron lactate shuttling to age-related changes in *D. melanogaster* memory has not been investigated.

Lactate dehydrogenase (LDH) is a tetrameric enzyme which catalyzes the interconversion of lactate and pyruvate coupled with the exchange of the oxidated and reduced forms of the coenzyme nicotinamide adenine dinucleotide (NAD) [Pyruvate + NADH + H^+^ ↔ Lactate + NAD^+^]. This reaction serves as a central node for balancing potential outcomes of carbohydrate metabolism. The breakdown of circulating sugars such as glucose, or trehalose in the case of *D. melanogaster,* is biased towards the production of pyruvate, which can be further processed by different metabolic enzymes [55]. Pyruvate can be oxidized by the pyruvate dehydrogenase complex to acetyl coenzyme A (Acetyl-CoA), followed by subsequent oxidization into carbon dioxide (CO2) through the tricarboxylic acid (TCA) cycle in mitochondria. Alternatively, pyruvate can be reduced to lactate by LDH and serve as a reservoir for oxidative metabolism later following the conversion back into pyruvate by LDH [56]. Therefore, glia rely on LDH to generate lactate, which is subsequently used as a fuel source by neurons which use LDH to convert lactate to pyruvate to fuel oxidative phosphorylation. Vertebrates have two isoforms of LDH expressed in the brain, LDHA and LDHB, which are believed to be differentially expressed in glia and neurons [57,58]. Five different LDH isozymes can be formed depending on which vertebrate LDH isoforms make up the subunit composition of the LDH tetramer (LDH-1 = B4, LDH-2 = B1A3, LDH-3 = B2A2, LDH-4 = B1A3, LDH-5 = A4). The enzyme kinetics of LDH isozymes differ such that those containing more LDHA favor conversion of pyruvate to lactate and those containing more LDHB favor the conversion of lactate to pyruvate [59–68]. In contrast, *D. melanogaster* LDH is a tetramer made up of a four identical subunits encoded by a single LDH homolog gene *ImpL3/dLdh* (FBgn0001258) [69,70] that is expressed in glia and neurons [71,72]. Interestingly, in neuronal nuclei of the mushroom body brain region, the main olfactory memory centre within *D. melanogaster, dLdh* mRNA levels are nearly 8-fold higher compared to the rest of the brain [72]. Moreover, aging in *D. melanogaster* causes an increase in *dLdh* mRNA [51,73–75] and dLdh protein levels in the brain [51]. In addition, adult specific *dLdh* overexpression, either ubiquitously or in a tissue specific manner in muscle, neurons, and glia, has recently been shown to decrease lifespan [51,76]. Nonetheless, the impact of altered *dLdh* expression in specific CNS cell types on memory, in an aging context, has not yet been examined in flies.

In *D. melanogaster,* age-related decline in cognitive function has been well documented [77,78]. One of the most utilized classical conditioning assays for assessing *D. melanogaster* learning and memory is the aversive olfactory associative learning task that requires exposing flies to two separate odors; one paired with electric shocks and the other without, then measuring preference for the non-paired odor between the two [79]. Olfactory associative memory is particularly sensitive to aging [80,81] and requires various types of metabolism depending on the training and retrieval conditions [48–50,82,83]. Another well characterized assay for assessing *D. melanogaster* learning and memory is the courtship conditioning paradigm. Courtship conditioning occurs when rejection by females of male courtship attempts promotes a reduction of future male courtship attempts [84]. Courtship memory is similar to aversive olfactory associative memory in that both behaviours require olfaction, depend on the mushroom body region of the brain, and can be retained for days upon appropriate training conditions [85–87].

Here we studied the effect of genetic alteration of *dLdh* expression within adult neurons or glia on long-term courtship memory and aging. We first show that long-term courtship memory is unaffected with age in our control flies (Canton-S); a finding in agreement with previous studies of short-term courtship memory [88] and in contrast to previous studies assessing olfactory based long-term memory [80,81]. However, we found that genetic changes in lactate metabolism result in alterations in long-term courtship memory in an age- and cell-type-dependent manner. We provide evidence that both increasing and decreasing *dLdh* expression within neurons negatively affects survival and maintenance of long-term courtship memory with age. Paradoxically, excess glial *dLdh* expression reduces lifespan without a change in memory, whereas decreased glial *dLdh* expression causes an age-dependent deficit in long-term courtship memory in the absence of any effect on lifespan. Furthermore, we show that changes in metabolite levels and neutral lipid accumulation may underlie age-related impacts of altered lactate metabolism, suggesting that altered glia-neuron lactate and lipid shuttling contribute to cognitive aging and decreased brain health in *D. melanogaster.*

## Results

### Lactate dehydrogenase expression and long-term courtship memory is maintained with age in Canton-S flies

In this study, courtship conditioning was employed to assess long-term memory (Figure 1A). To date, there are no reports examining the effects of age on long-term courtship memory in a reference or control line such as Canton-S (CS). We observed that 24-hour long-term courtship memory was retained in CS up to 30 days of age under standard growth temperature conditions (25°C) (Figure 1B + 1C). These results indicate that long-term courtship memory is not susceptible to age-related impairment, at least up to 30 days of age at 25°C, a time point when 89.7% of the population is still alive.

**Figure 1.**
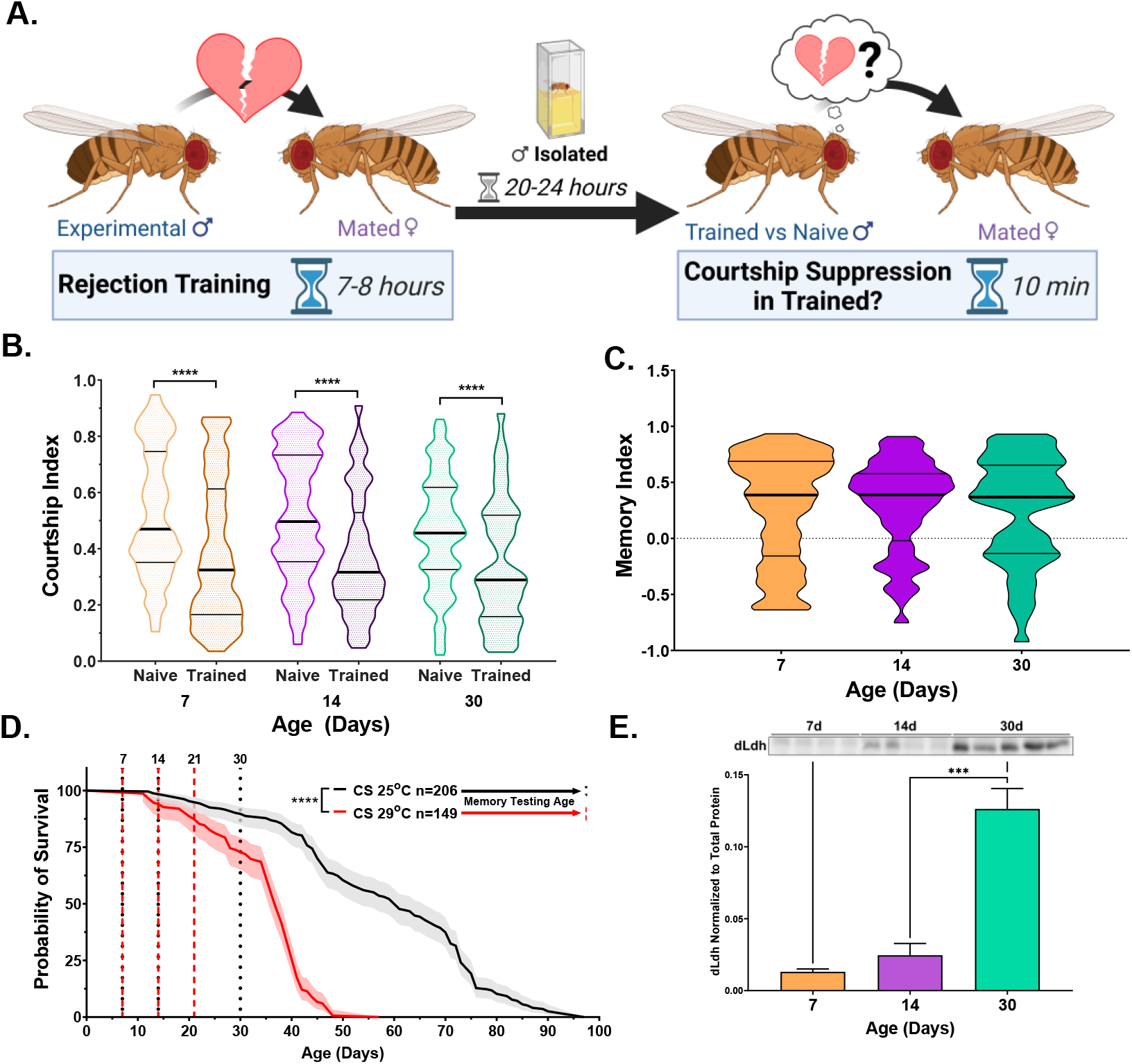
dLdh protein levels increase in the brain with age while long-term courtship memory is retained in control flies. **A.** Courtship conditioning paradigm used for testing long-term courtship memory throughout this study. **B**. Courtship indexes for Canton-S male flies aged 7, 14, or 30 days at 25°C decreased in courtship conditioning rejection trained vs naïve conditions at all ages. N=100-117. Naïve and trained flies were compared for each age group using one-sided Mann-Whitney U tests, **** = *p*<0.0001. Data presented as violin plot of frequency distribution. **C.** Long-term courtship memory indexes for trained flies represented in B do not differ between age groups. Age groups were compared using a Kruskal-Wallis test with Dunn’s multiple comparisons tests. **D.** Survival curve of Canton-S male flies singly housed is decreased at 29°C compared with 25°C and ages which were selected for memory testing in control (25°C black) and transgenic (29°C red) flies are highlighted with vertical dotted lines. Curve comparison was made using a Log-rank (Mantel-Cox) test, **** = *p*<0.0001. Data presented as violin plot of frequency distribution. **E.** Western blot analysis of head extracts from male Canton-S flies aged 7, 14 or 30 days at 29°C showing dLdh protein levels increase with age (*F*(2, 10)=36.64, *p*<0.0001) n=4-5. Each sample consists of protein extracts from 20 heads. Comparisons across age were made using one-way ANOVA with Šídák’s multiple comparisons tests between all age groups. *** = *p*<0.001, **** = *p*<0.0001.

We restricted genetic manipulation of *dLdh* expression to adulthood by employing a system which utilizes temperature sensitive Gal80 to inhibit Gal4 activity at 18°C, while permitting activity at 29°C [89]. We set up a reference longevity curve, in which CS flies were reared and assayed at 29°C. At this temperature, flies exhibited approximately 90% survival at 21 days of age, which is similar to CS flies reared at 25°C at 30 days of age (Figure 1D). Thus, to limit survivorship bias we selected 21 days of age raised at 29°C for analysis of aged flies. We also examined endogenous levels of dLdh at this timepoint by western blot analysis. CS flies reared at 29°C exhibited an increase in dLdh protein levels with age (Figure 1E), in accordance with our previous evaluation of aged cantonized *w^1118^* flies maintained at 25°C [51] and with others who have measured *dLdh* only at the transcript level [73–75,90].

### Both increasing and decreasing neuronal lactate dehydrogenase expression is detrimental to survival and long-term courtship memory in aged flies

In order to assess the effects of altered *dLdh* expression on memory in an adult context, we used flies containing a temperature sensitive Gal80 driven by a ubiquitously expressing Tubulin driver and Gal4 driven by a pan-neuronal expressing Elav driver (thereafter Elavts-Gal4). These flies were subsequently crossed with flies containing an upstream activation sequence (UAS) driving either the dLdh full-length open reading frame with a triple hemagglutinin (HA) tag on the C-terminus (UAS-dLdh) or an RNA interference construct targeting *dLdh* (UAS-dLdh-RNAi), to permit either increased or decreased expression of *dLdh* in adult neurons respectively. Firstly, we verified by western blot that dLdh protein levels decreased in the heads of neuronal *dLdh* downregulated flies and that there was induction of HA-tagged dLdh production in the heads of neuronal *dLdh* upregulated flies at 21 days of age (Figure 2A). Using the same courtship conditioning paradigm utilized in CS flies, we tested long-term courtship memory in dLdh transgenic flies. At 7 and 14 days of age, neither neuronal *dLdh* upregulation nor downregulation elicited a change in long-term courtship memory (Figure 2B + 2C). However, at 21 days of age, flies with either a neuronal upregulation or downregulation of *dLdh* expression showed a decrease in long-term courtship memory compared to control flies (Figure 2B + 2C). In addition to memory, we tested whether aging affected health in neuronal *dLdh* transgenic flies by measuring survival and climbing ability. Flies with neuronal *dLdh* downregulation exhibited a clear decrease in survival compared to control flies, with an earlier onset of death, a pronounced increase in the rate of death, but an unchanged maximum lifespan (Figure 2D). Flies with neuronal *dLdh* upregulation also exhibited reduced survival compared to control flies, with a similar decrease in time of death onset and rate of death, but also with a decrease in maximum lifespan (Figure 2D). Unhealthy flies tend to exhibit a general inability to maintain locomotor behaviours as assessed using a negative geotaxis assay. We found that climbing ability was decreased in flies with neuronal *dLdh* upregulation compared to control flies at 21 days of age (Figure 2E). These findings indicate that the rate of age-related decline in climbing ability of flies with neuronal *dLdh* upregulation was more pronounced than control flies. However, flies with neuronal *dLdh* downregulation did not exhibit a change in climbing ability compared to control flies at any age (Figure 2E).

**Figure 2.**
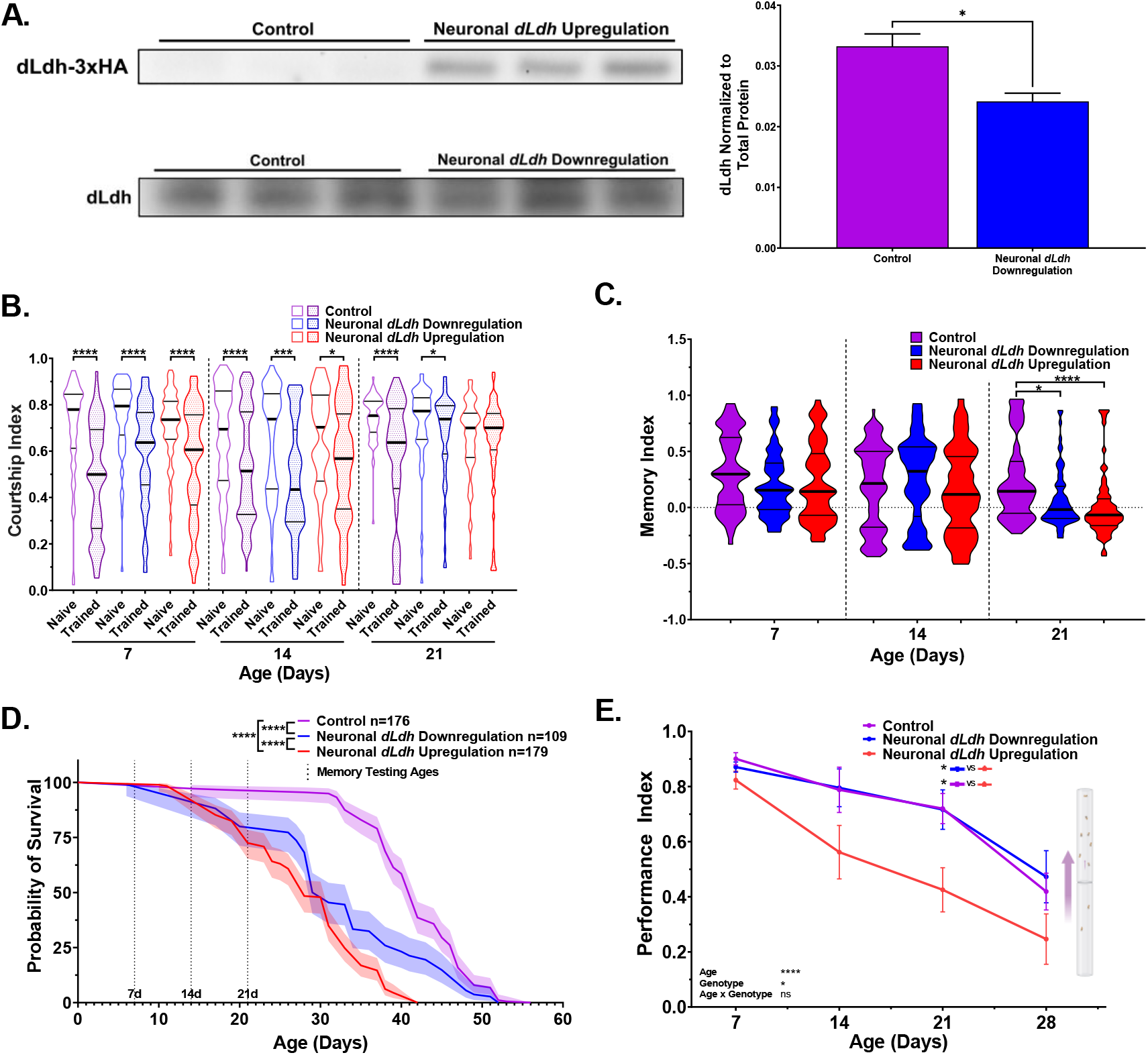
Flies with either elevated or downregulated neuronal *dLdh* expression exhibit decreased long-term courtship memory and survival with age. **A.** Western blot analysis of head extracts from neuronal transgenic male flies aged 21 days at 29°C showing elevated ectopic HA tagged dLdh expression (upper panel) and decreased endogenous dLdh levels (lower panel) in flies using a Elavts-Gal4 driver to drive UAS-dLDH (with 3xHA tag) and UAS-dLdh-RNAi expression respectively (n=3), expression was normalized to total protein content (based on Ponceau-S staining). Each sample consisted of protein extracts from 20 heads. Comparison of endogenous dLdh level were made using an unpaired t test, * = *p*<0.05. **B.** Courtship indexes for neuronal transgenic male flies aged 7, 14, or 21 days at 29°C post-eclosion showed a decrease in courtship conditioning rejection trained vs naïve for all genotypes at all ages, except for 21 day upregulation. N=59-125. Naïve and trained flies were compared within each genotype at each age using one-sided Mann-Whitney U tests, ****=*p*<0.0001, ***=*p*<0.001, *=p<0.05. Data presented as violin plot of frequency distribution. **C.** Long-term courtship memory indexes for neuronal transgenic trained flies represented in B. Memory in neuronal transgenic flies only differed from control at 21 days of age (*H*=21.33, *p*<0.0001), with both flies with *dLdh* upregulation and downregulation showing reduced memory at 21 day of age compared to control. Genotypes at each age were compared using Kruskal-Wallis tests with Dunn’s multiple comparisons to control, ****=*p*<0.0001, *=*p*<0.05. Data presented as violin plot of frequency distribution. **D.** Both neuronal upregulation and downregulation of *dLdh* resulted *in* reduced survival compared to control. Shaded area represents the 95% confidence interval. Curve comparisons were made using a Log-rank (Mantel-Cox) test, **** = *p*<0.0001. Ages chosen for memory testing are highlighted with vertical dotted lines. **E.** Upregulation of neuronal *dLdh* decreased climbing ability during negative geotaxis compared to control and *dLdh* downregulation (genotype effect *F*(2, 33)=4.355, *p*=0.0210). Comparisons across age and genotype were made using a mixed-effects model with Geisser-Greenhouse correction and Šídák’s multiple comparisons within each age group, * = *p*<0.05, ****=p<0.0001. Effects of age, genotype, and age by genotype interaction are denoted on the bottom left.

### Altered glial lactate dehydrogenase expression promotes differential effects on survival and long-term courtship memory

To restrict manipulation of *dLdh* expression to glial cells in adults, we employed flies containing a pan-glial expressing Repo driver combined with the same temperature system used for neuronal *dLdh* overexpression (Repots-Gal4). We verified decreased dLdh protein levels in the heads of glial *dLdh* downregulated flies and induced HA-tagged dLdh levels in the heads of glial *dLdh* upregulated flies at 21 days of age by western blot analysis (Figure 3A). We used courtship conditioning to assess memory in these flies. Glial *dLdh* transgenic flies exhibited no significant differences in memory compared to control flies at 7 and 14 days of age (Figure 3B + 3C), indicating that long-term courtship memory was unimpacted by glial *dLdh* manipulation at these ages. In contrast, a clear deficit in long-term courtship memory was observed in flies with glial *dLdh* downregulation compared to control flies at 21 days of age (Figure 3B + 3C). Glial *dLdh* upregulation had no effect on memory at 21 days of age (Figure 3B + 3C). The effect of altered glial *dLdh* expression on lifespan was also assessed. We detected a reduction in the survival of flies with glial *dLdh* upregulation compared to control flies, with an earlier onset of death, a slight increase in the rate of death, and minimal change in maximum lifespan (Figure 3D). Flies with glial *dLdh* downregulation had indistinguishable survival compared to control flies (Figure 3D). The climbing ability of flies with either glial *dLdh* upregulation or downregulation did not differ from control flies at all ages (Figure 3E).

**Figure 3.**
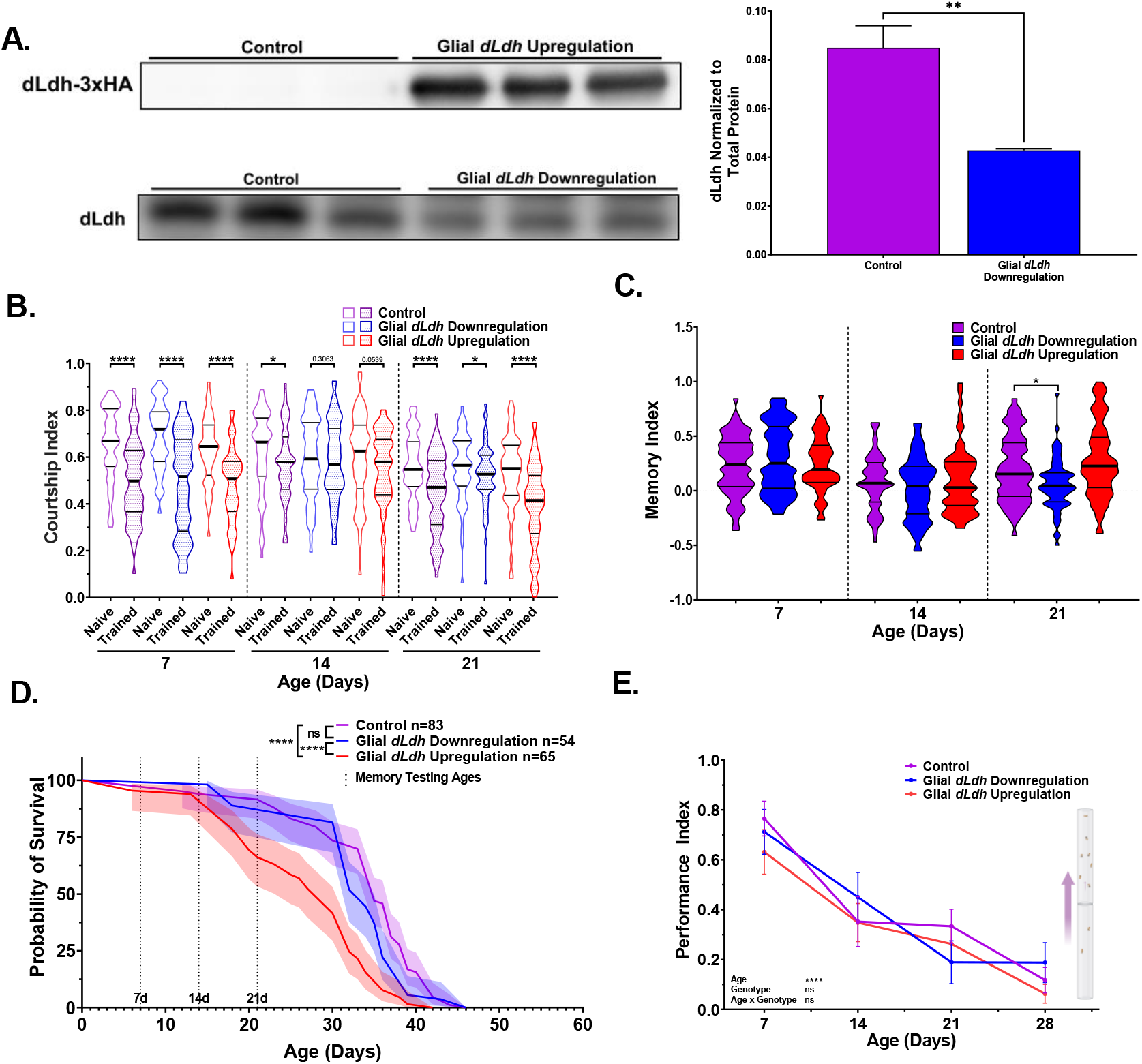
Flies with downregulated *dLdh* expression in glia exhibit decreased long-term courtship memory with age, while flies with upregulated *dLdh* expression in glia exhibit decreased survival. **A.** Western blot analysis of head extracts from glial transgenic male flies aged 21 days at 29°C showing elevated ectopic HA tagged dLdh expression (upper panel) and decreased endogenous dLdh levels (lower panel) in flies using a Repots-Gal4 driver to drive UAS-dLDH (with 3xHA tag) and UAS-dLdh-RNAi expression respectively (n=3), expression was normalized to total protein content (based on Ponceau-S staining) n=3. Each sample consisted of head extracts from 20 heads. Comparisons made by unpaired t test, ** = *p* < 0.01 **B.** Courtship indexes for glial transgenic male flies aged 7, 14, or 21 days at 29°C post-eclosion showing courtship conditioning rejection training was decreased compared to naïve flies for all genotypes at 7 and 21 days of age, and control only at 14 days of age. n=48-72. Naïve and trained flies were compared within each genotype at each age using one-sided Mann-Whitney U tests, ****=*p*<0.0001, *=*p*<0.05. Data presented as violin plot of frequency distribution. **C.** Long-term courtship memory indexes for glial transgenic trained flies represented in B. Memory was only decreased following glial *dLdh* downregulation at 21 days of age compared to control (*H*=16.9*, p*=0.0002). Genotypes at each age were compared using Kruskal-Wallis tests with Dunn’s multiple comparisons to control, *=*p*<0.05. Data presented as violin plot of frequency distribution. **D.** Only glial upregulation of *dLdh* reduced survival compared to control and *dLdh* downregulation. Shaded area represents the 95% confidence interval. Curve comparison was made using a Log-rank (Mantel-Cox) test, **** = *p* < 0.0001. Ages chosen for memory testing are highlighted with vertical dotted lines. **E.** Altered glial *dLdh* expression did not impact climbing ability during negative geotaxis at any age compared to control. Comparisons across age and genotype were made using a mixed-effects model with Geisser-Greenhouse correction and Tukey’s multiple comparisons within each age group. Effects of age, genotype, and age by genotype interaction are denoted on the bottom left.

### Elevated *dLdh* expression in either neurons or glia promotes neutral lipid accumulation in the brains of aged flies

Glial generated lactate is a substrate for neuronal lipid synthesis [36]. Therefore, we investigated whether lipids were altered in the brains of aged transgenic flies in which either neuronal or glial lactate metabolism was genetically manipulated. We used nile red staining on whole brains dissected from flies at 21 days of age to detect neutral lipids [91,92]. Brains of flies with *dLdh* upregulated in either neurons or in glia displayed a pronounced increase in neutral lipids compared to control flies (Figure 4). Downregulation of *dLdh* in either neurons or glia had no apparent impact on neutral lipids detected in the brain, as assessed by nile red staining (Figure 4).

**Figure 4.**
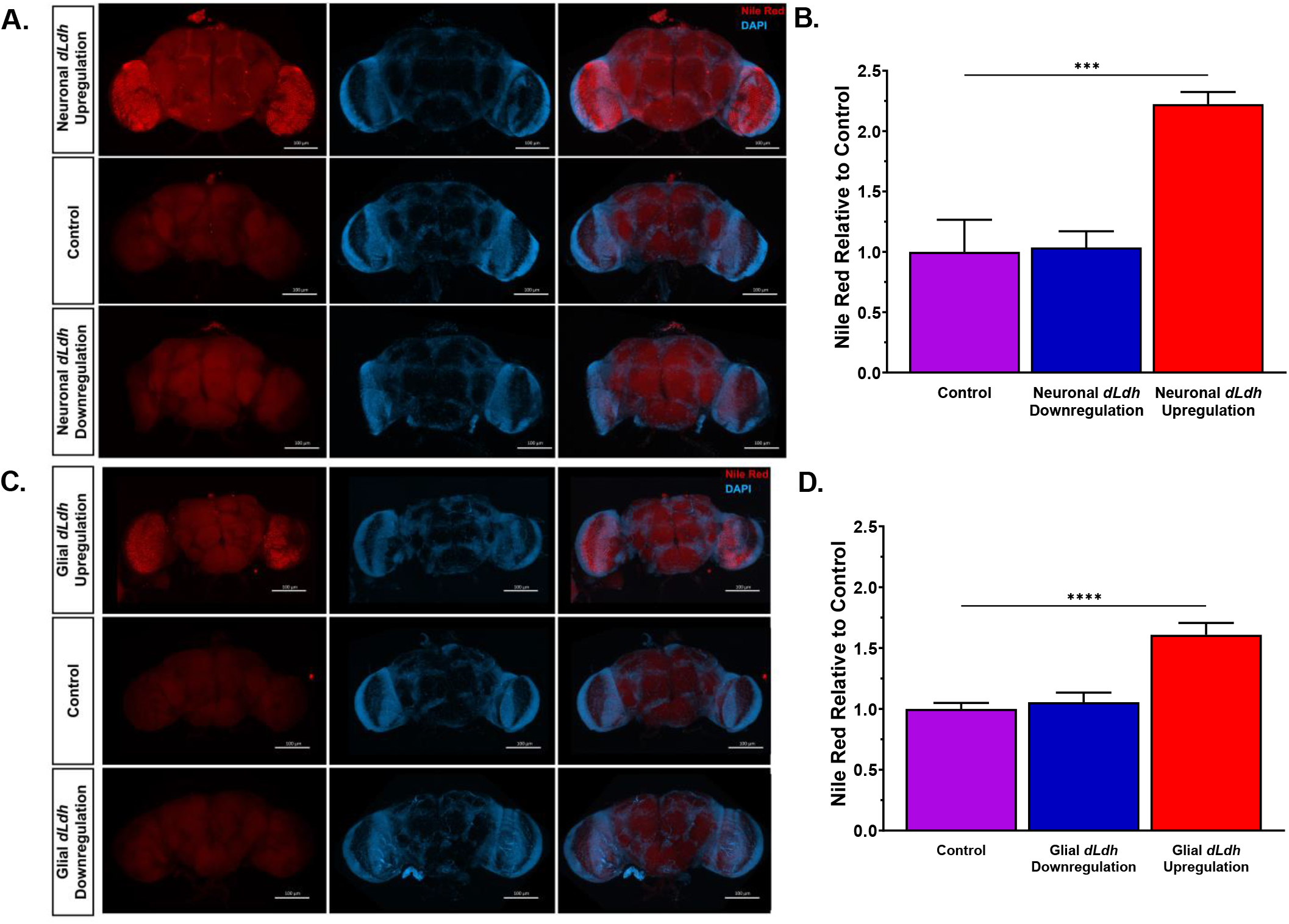
Increased glial or neuronal *dLdh* expression promotes accumulation of neutral lipids in 21 day aged male transgenic fly whole brains. **A.** Representative confocal fluorescence microscopy images of neuronal *dLdh* transgenic fly brains removed at 21 days of age at 29°C and stained for DNA (Blue = DAPI) and neutral lipids (Red = Nile Red). **B.** Quantification nile red staining mean fluorescence intensity revealed that flies with neuronal *dLdh* upregulation have over twice the lipid accumulation as control flies. N=4-5 (*F*(2, 10) = 17.69, *p*=0.0005) **C.** Representative confocal fluorescence microscopy images of glial *dLdh* transgenic fly brains dissected out at 21 days of age and stained for DNA (Blue = DAPI) and neutral lipids (Red = Nile Red). **D.** Quantification nile red staining mean fluorescence intensity revealed that flies with glial *dLdh* upregulation have over 1.5 times lipid accumulation as control flies. N=7-8. (*F*(2, 19) = 19.74, *p*<0.0001). Comparisons between genotypes for neuronal and glial *dLdh* transgenic flies separately were made using one-way ANOVA with Dunnett’s multiple comparisons with control. *** = *p*<0.001, **** = *p*<0.0001.

### Brain metabolite levels in the heads of flies fluctuate with age and are readily altered by manipulating glial or neuronal lactate dehydrogenase expression

To determine how manipulation of *dLdh* in neurons or glia impact age-related changes in metabolism across the brain, we assessed metabolites extracted from transgenic fly heads at 7 and 21 days of age using gas chromatography-mass spectrometry (Figure 5, Figure 5S). We initially focused on lactate and pyruvate as these are the metabolites that are directly impacted by dLdh canonical activity (Figure 5A). Surprisingly, brain levels of lactate were unaffected by neuronal *dLdh* manipulation (Figure 5B).

**Figure 5.**
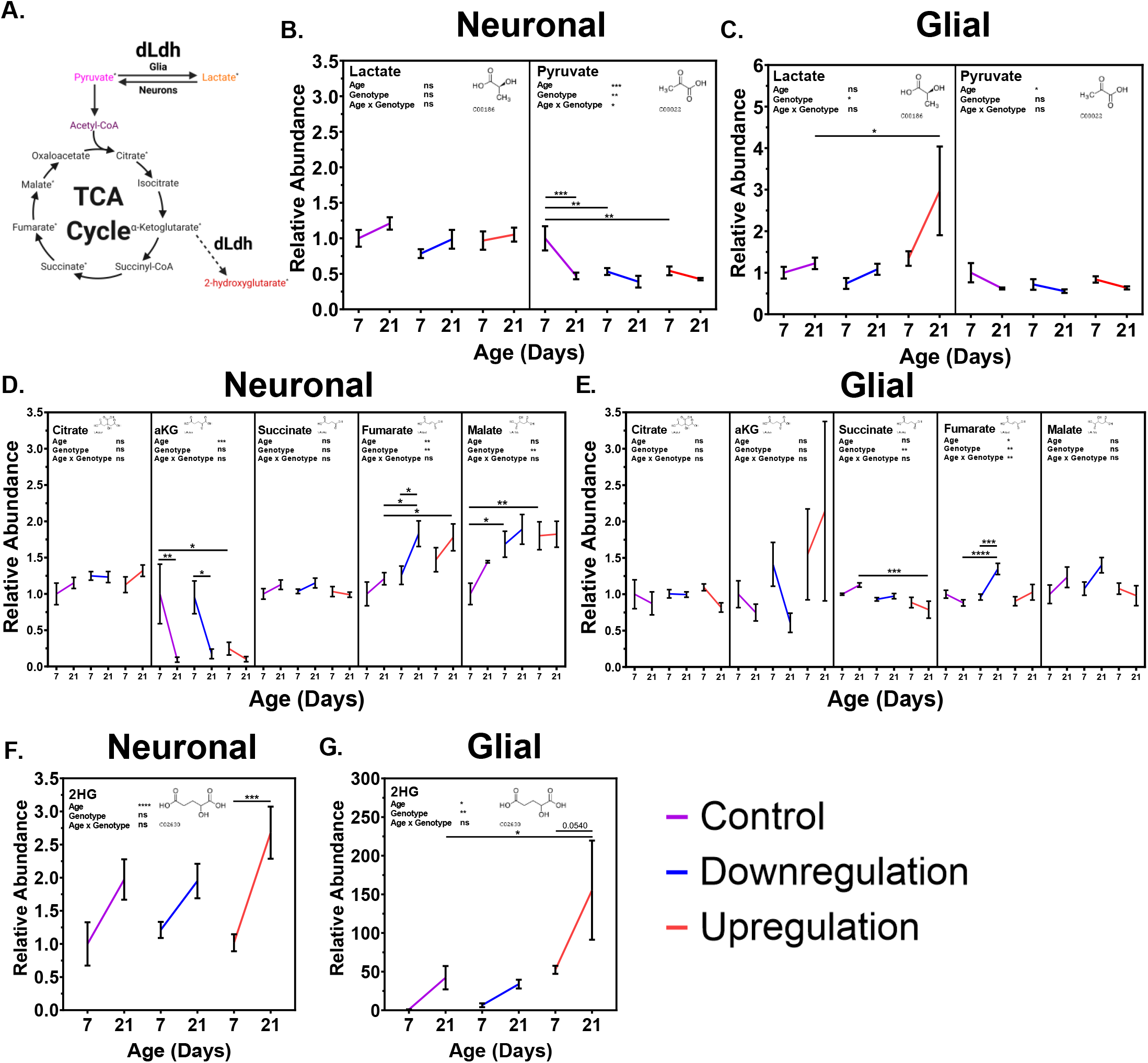
Metabolite analysis of transgenic male fly heads aged 7 and 21 days with altered expression of neuronal or glial *dLdh* revealed alterations in lactate, pyruvate, TCA cycle intermediates, and 2HG levels relative to control flies. **A.** Metabolic pathway connections with metabolites produced by dLdh canonical [Pyruvate ↔ Lactate] and non-canonical [αKG→2HG] activity. Metabolites measured by gas chromatography-mass spectrometry (GC-MS) in the heads of transgenic male flies aged 7 and 21 days at 29°C are denoted by a superscript asterisk. **B.** Neuronal *dLdh* manipulation does not impact lactate levels whereas pyruvate levels are reduced at 7 days of age nearly to the level of 21 days of age by both upregulation and downregulation. **C.** Glial *dLdh* upregulation caused an increase in lactate but no effect on pyruvate levels. **D.** Neuronal *dLdh* manipulation promotes alterations in select TCA cycle intermediates. Upregulation of *dLdh* lowered α-ketoglutarate (αKG) levels. Downregulation and upregulation of *dLdh* raised fumarate and malate levels. **E.** Glial *dLdh* manipulation caused only slight alterations in TCA cycle intermediates. Citrate, αKG, and malate levels were unaltered. Upregulation of *dLdh* lowered succinate levels. Downregulation of *dLdh* raised fumarate levels at 21 days of age. **F.** 2-hydroxyglutarate (2HG) levels are generally increased with age without any clear change due to neuronal dLdh manipulation. **G.** The age-related increase in 2HG is exacerbated by glial *dLdh* upregulation but not by downregulation. Comparisons for each metabolite were done between genotype and age groups of neuronal and glial *dLdh* transgenic flies separately using two-way ANOVAs with Dunnet’s multiple comparisons with control for each age group and with Šídák’s multiple comparisons between age groups within each genotype. Effects of age, genotype, and age by genotype interaction are denoted on the top left of each graph. The structural formula for each metabolite were obtained from the Kyoto Encyclopedia of Genes and Genomes (KEGG) chemical compound database with associated C number below.

However, the age-related decline in brain pyruvate evident in control flies was prematurely triggered by both downregulation and upregulation of neuronal *dLdh,* with a clear reduction at 7 days of age that was maintained until 21 days of age in neuronal transgenic flies (Figure 5B). Glial *dLdh* upregulation caused a pronounced accumulation of lactate at 21 days of age, whereas downregulation had no impact on overall brain lactate levels (Figure 5C). The age-related decline in brain pyruvate levels was unaffected by both glial *dLdh* upregulation and downregulation (Figure 5C). Next, we examined the effects of altered *dLdh* expression on mitochondrial metabolism by assessing changes in TCA cycle intermediates, including citrate, alpha-ketoglutarate (αKG), succinate, fumarate, and malate (Figure 5A). We saw no differences in brain citrate level regardless of age or *dLdh* manipulation (Figure 5D + 5E). In control flies for neuronal *dLdh* manipulation, brain levels of αKG showed a substantial decline from 7 to 21 days of age (Figure 5D), whereas in neuronal *dLdh* upregulation flies, low levels of αKG in the brain were detected at both 7 and 21 days of age (Figure 5D). In both the glial *dLdh* upregulation and downregulation lines there were no changes in brain αKG levels at either 7 or 21 days of age (Figure 5E). A small decrease in brain succinate levels was observed in flies with glial *dLdh* upregulation compared to the control flies, particularly at 21 days of age (Figure 5E). Both upregulation and downregulation of neuronal *dLdh* promoted an age-related increase in brain fumarate levels compared with the control flies (Figure 5D). An age-related elevation in brain fumarate was only evident in flies with glial *dLdh* downregulation but not upregulation (Figure 5E). Malate levels in the brain were increased in both neuronal *dLdh* upregulated and downregulated flies, but this difference was most pronounced at 7 days of age relative to control flies (Figure 5D). Glial *dLdh* manipulation had no effect on malate levels at any age (Figure 5E). Lastly, we measured brain levels of 2-hydroxyglutarate (2HG), which consists of both the D- and L-enantiomers (Figure 5A). In both control and neuronal *dLdh* transgenic flies we detected increased 2HG levels in brains with age. (Figure 5F). Interestingly, we found brain 2HG levels to be increased in flies with glial *dLdh* upregulation compared with control, with approximately 155-fold higher 2HG levels at 21 days of age compared to 7-day old control flies (Figure 5G). Considering that dLdh catalyzes the formation of L-2HG from αKG, this pronounced increase in 2HG level likely represents increased production of L-2HG.

## Discussion

### dLdh expression needs to be precisely regulated to ensure optimal aging

Here we show that downregulation of *dLdh* in neurons, but not glia, leads to lower survival. In accordance with glia-neuron lactate shuttling, the lower survival caused by neuronal downregulation of *dLdh* may be attributed to the decreased ability of neurons to oxidize glial-derived lactate to pyruvate in order to fuel synaptic activity and neuronal maintenance. Indeed, we detected decreased pyruvate levels in neuronal dLdh downregulated flies at 7 days of age compared to control flies. A previous study demonstrated that suppressing glycolytic enzyme expression in glia resulted in the decreased ability of glia to process trehalose as a fuel source, with a corresponding reduction in survival [34]. We previously demonstrated that glial *dLdh* downregulation in group housed flies led to increased survival [51]. However, in the current study, we detected no change in survival in males with glial *dLdh* downregulation, reared in isolation. Social isolation is known to impact aging, as well as many physiological processes [93–95]. It is possible that interfering with the ability of neurons to oxidize lactate in neuronal *dLdh* downregulation flies prevents neurons from freely shifting their metabolism in an adaptive manner to accommodate age-related changes in nutrient availability and mitochondrial oxidative capacity. In contrast, decreased glial-generated lactate in glial *dLdh* downregulation flies still allows neurons to utilize cell-autonomously generated lactate or to adapt by shifting metabolism towards utilization of other types of fuel sources or employing alternative metabolic pathways.

The question arises as to why survival was not decreased by glial *dLdh* downregulation in a manner similar to that others have observed following downregulation of other glycolytic enzymes [34]? We propose that a lack of glial dLdh-derived lactate for fueling neurons could be balanced by preventing damage from excess dLdh activity arising with age. The adverse effects elicited by elevated dLdh activity are exemplified by our results here showing that increasing *dLdh* in either glia or neurons causes decreased survival in *D. melanogaster.* In a previous study we found that neuronal *dLdh* upregulation caused an increase in the number of brain vacuoles [51], indicative of neurodegeneration occurring in the fly brain with age [96]. Other studies in flies have recently shown that the toxic effect of amyloid beta, a peptide strongly implicated in neurodegeneration in Alzheimer’s disease, causes *dLdh* upregulation in part due to an Activating transcription factor 4 (ATF-4) dependent endoplasmic reticulum stress signalling unfolded protein response [52]. In accordance with our findings on neuronal dLdh impact on fly survival, the authors of this study also found that both upregulation and downregulation of neuronal *dLdh* caused an exacerbation of amyloid beta toxicity [52]. Furthermore, a recent finding demonstrated that ubiquitous disruption of the *Ldhb* gene within mice resulted in increased mitochondrial dysfunction, oxidative stress, apoptosis, and cognitive impairment [97]. However, this study did not distinguish which cell type was adversely affected nor did it assess the effects of reduced *Ldhb* expression on global metabolism and brain health with age. It will be essential for future studies to determine if altered lactate metabolism in flies promotes neuronal loss via programed cell death pathways such as in apoptosis, necroptosis, or ferroptosis with age.

### dLdh impacts age-related changes in brain metabolism differentially in neurons and glia

Aging in flies is associated with higher TCA cycle activity and nicotinamide adenine dinucleotide + (NAD+) consumption [98], in addition to a decline in mitochondrial electron transport chain (ETC) activity [74,99,100]. One possible explanation for decreased survival in neuronal *dLdh* overexpression is that chronic lactate oxidation in neurons may have triggered an age-related shift in neuronal metabolism towards accelerated TCA cycle activity at a younger age and diminished ability of neurons to slowly adapt to other age-related changes. The metabolic profile of neuronal *dLdh* transgenic flies in this study supports this claim. Pyruvate feeds the TCA cycle following conversion to acetyl-CoA and the subsequent formation of a succession of intermediates including citrate, isocitrate, αKG, succinyl-CoA, succinate, fumarate, malate, and ends with oxaloacetate that can react with a new acetyl-CoA to start the cycle again. We found a premature decrease in pyruvate in the heads of neuronal *dLdh* upregulated flies which may have arisen due to neurons processing pyruvate more readily through the TCA cycle at an earlier age. Moreover, in neuronal *dLdh* upregulation fly heads we saw a premature decrease in αKG, which is produced earlier in the TCA cycle after entry of pyruvate via acetyl-CoA, along with a premature increase in fumarate and malate, which are metabolites later in the TCA cycle relative to pyruvate. These changes in TCA cycle intermediates could be an indication of a higher propensity for neuronal TCA cycle activity in flies with neuronal *dLdh* upregulation.

Similarly in neuronal *dLdh* downregulated flies we saw a premature decrease in pyruvate and an increase in TCA cycle intermediates further downstream of pyruvate, including fumarate and malate, yet αKG was unchanged. If neurons with downregulated *dLdh* have reduced ability to oxidize lactate, then an increased TCA cycle activity in neurons could be due to increased provision of non-lactate derived pyruvate sources or an increase in metabolites entering the TCA cycle through anaplerotic reactions. For example, serine is an amino acid which can function as a source of pyruvate when acted on by enzymes with L-serine ammonia-lyase activity. A gene (FBgn0037684) was recently identified in the *D. melanogaster* brain that encodes for a protein predicted to have this function [101,102]. We detected a particularly high level of serine in the heads of flies with neuronal *dLdh* downregulation or glial *dLdh* upregulation (Figure 5S). Studies in mice suggest serine can contribute to astrocyte metabolism [103], memory loss in Alzheimer’s disease [104], oxidative stress response in T cells [105], and fuel for one-carbon metabolism [106,107]. Moreover, D-serine is a co-agonist of N-methyl-D-aspartate (NMDA) receptors that is produced by both neurons and glia [108] and involved in age-related memory impairment of short-term aversive olfactory associative memory in flies [54]. Therefore, further experiments are required to elucidate how the role of serine is involved in the impact of lactate metabolism on the aging brain.

Elevated fumarate levels were detected in the heads of both neuronal *dLdh* upregulation and downregulation flies which could be due to elevated succinate dehydrogenase (SDH) activity in the TCA cycle and ETC. SDH is the only complex in the ETC which is not encoded by genes in the mitochondrial genome. Therefore, SDH function may be less impacted with age than other ETC complexes because the genes encoding the SDH complex are not as susceptible to age-related increases in oxidative damage compared to mitochondrial genes [109–111]. Overall, our metabolite analysis suggests that *D. melanogaster* neurons require finely regulated expression of *dLdh* to cope with age-related brain metabolic changes, such as high TCA cycle activity and mitochondrial dysfunction, to ensure optimal lifespan.

Although flies with both glial and neuronal dLdh upregulation exhibited shortened lifespans, the survival curves for both lines revealed subtle differences. While neuronal and glial *dLdh* upregulated flies had an increase in the rate of death and an earlier onset of death, only neuronal *dLdh* upregulated flies had a reduced maximum lifespan. In addition, neuronal *dLdh* upregulated flies were the only *dLdh* manipulated group which showed a decrease in climbing ability compared to control flies, indicative of a lower healthspan [112]. An earlier onset of aging, with no apparent effect on the mortality rate, could be responsible for the decrease in maximum longevity we observed when *dLdh* was upregulated in neurons. In contrast, an early onset of aging, associated with a decreased mortality rate, best explains the unaffected maximum lifespan we saw in flies with *dLdh* upregulated in glia. Therefore, an age-related increase in neuronal TCA cycle activity may not necessarily explain these glial *dLdh* induced survival deficits. This is corroborated by our observation that TCA cycle intermediates were unchanged in the heads of glial *dLdh* transgenic flies. The only change in TCA cycle intermediates that we saw in flies with glial *dLdh* upregulation was a slight drop in succinate. Interestingly, we detected an accumulation of lactate only in the heads of flies with glial but not neuronal *dLdh* upregulation. Because we were only able to detect pyruvate changes in heads of neuronal transgenic flies and lactate changes in heads of glial transgenic flies, these observations suggests that in *D. melanogaster* heads it is likely that neurons dictate pyruvate levels and glia are more responsible for lactate levels. Moreover, *D. melanogaster* neurons may lack the ability to release lactate, unlike glia [34]. Therefore, glia with *dLdh* upregulation may produce excess lactate above the capacity of neurons to uptake and oxidize it, causing lactate to accumulate to detrimental levels. Overall, our metabolite analysis suggests that decreased survival in glial *dLdh* upregulation files cannot be fully explained by age-related changes in TCA cycle intermediates, but glial overproduction of lactate may be a contributing factor.

### L-2HG may be a novel biomarker of brain aging which is increased by glial dLdh

Another metabolite which could be relevant to survival deficits caused by altered *dLdh* expression and aging is L-2HG. In the heads of flies with *dLdh* altered in neurons and their respective genotype control, we found a sharp increase in 2HG with age. Interestingly, we found an even larger increase of 2HG with age in the heads of flies with *dLdh* altered in glia and their respective genotype control; with 2HG elevated 155-fold by *dLdh* upregulation compared to control. L-2HG is a common product of Dipteran larval metabolism [113]. L-2HG accumulates in *D. melanogaster* larvae due to non-canonical activity of dLdh leading to the conversion of αKG to L-2HG [114], or by inhibition the enzyme L-2-hydroxyglutarate dehydrogenase (dL2HGDH) that degrades L-2HG [114]. However, L-2HG levels in *D. melanogaster* are much lower in adults compared to larvae [114] and a recent study searching for biomarkers of aging in *D. melanogaster* did not even measure 2HG [115]. To our knowledge, we present here the first demonstration that 2HG increases in the brain of flies with age. One caveat with these findings is that our method of 2HG detection included both L-2HG and D-2HG. Although D- and L-2HG are present at similar levels in adults [113], the two enantiomers are produced by distinct enzymatic mechanisms. Since *dLdh* expression is elevated in aging brains and glial *dLdh* upregulation drives a dramatic increase in 2HG levels, our findings are likely attributable to L-2HG accumulation. In the past, L-2HG has only been found to increase to appreciable levels in the adult brain of *D. melanogaster* through an ATF4-dependent unfolded protein response when mitochondrial stress is induced [116], or in flies exposed to hypoxic conditions [113]. Therefore, aging may cause accumulation of L-2HG due to a change in brain metabolism reminiscent of responses to mitochondrial stress or hypoxia, and these metabolic pathways may be further activated in the brains of glial *dLdh* upregulation flies due to elevated lactate accumulation. Furthermore, the lack of changes in lactate and L-2HG levels in neuronal *dLdh* upregulated fly heads compared to control flies suggests that dLdh may not be the sole contributor to lactate production in *D. melanogaster* neurons while elevated levels of L-2HG in older flies may require lactate-mediated inhibition of L-2HG degradation in addition to non-canonical dLdh activity generating L-2HG. While L-2HG may serve as a biomarker of an age-related shift in brain metabolism, elevated L-2HG levels may directly cause damage to the brain as well. L-2HG can directly inhibit ATP synthase (Complex I in the mitochondrial ETC) [117] and αKG dehydrogenase [118], provoke oxidative DNA damage [119], and cause severe neurological problems when it accumulates in children with inborn 2-hydroxyglutaric acidurias [120] or late-onset neurodegeneration when it accumulates in mice [121]. In fact, L-2HG production in response to age-related brain metabolic changes may be a general feature of brain aging and may be partially responsible for decreased survival caused by glial *dLdh* upregulation and lactate accumulation.

### Accumulation of neutral lipids in the brain may underlie age-related impacts of dLdh

There is an increasing recognition of the role that lipid metabolism plays in aging [122]. Lipids are typically stored as triglycerides and sterol esters in lipid droplet intracellular organelles for energy production [123,124]. Neutral lipids are covered in a monolayer of phospholipids and proteins in lipid droplets and can be broken down into fatty acid and cholesterol by lipolysis or autophagy (lipophagy) [125,126]. Here we show a marked increase in neutral lipids in the brain of aged flies with *dLdh* upregulated in either glia or neurons, suggesting lipid droplet accumulation in both cases. Accumulation of lipid droplets is known to occur in the brain during development [127,128], upon aging [129–131] and in response to stressors such as mitochondrial dysfunction [36,132], oxidative stress [91,132–134], ER stress [135,136], inflammation [131], starvation [91,136], and neurodegenerative disease [137–140]. Outside of lipid energy storage, there are a variety of potential lipid droplet functions, including protein maturation and turnover, intracellular motility, transcriptional regulation in the nucleus, and storage of proteins, vitamins, and signaling precursors [123]. In the brain, glia form lipid droplets more readily than neurons [125]. *D. melanogaster* larval brains accumulate lipid droplets exclusively in glia [127], whereas adulthood lipid droplets are predominantly localized in glia [91,133] and neurons to a lesser extent [141,142]. Neurons retain less lipid droplets than glia normally to avoid lipotoxicity occurring during the production of reactive oxygen species from free fatty acid β-oxidation [143,144]. Interestingly, it has been shown in mice and flies that the glia-neuron lactate shuttle provides a neuronal substrate for synthesis of fatty acids, but neurons avoid fatty acid lipotoxicity by shuttling fatty acids via apolipoproteins back to glia for production of lipid droplets [36,132,139,145–147]. The trading of lactate for fatty acids between glia and neurons to trigger glial lipid droplet production was shown to occur in the *D. melanogaster* retina and was prevented by knockdown of *dLdh* in either all neurons or pigment glia residing in the ommatidia [36]. Our results demonstrate that overproduction of *dLdh* in neurons or glia is sufficient to trigger neutral lipid accumulation outside of ommatidia and in the rest of the brain as well. Moreover, if neutral lipid accumulation is an indication of glial lipid droplets forming as an adaptive response to an age-related increase in neuronal mitochondrial dysfunction and oxidative stress, then elevation of these processes brain wide may partly explain why we observed reduced survival in flies with glial or neuronal *dLdh* upregulation in this study. In addition, if neutral lipid accumulation is an indication of increased neuronal lipid droplet formation, then survival deficits in flies with glial or neuronal *dLdh* upregulation could be due to a shift in neuronal metabolism towards excessive neuronal triglyceride synthesis which can trigger lipotoxicity when utilized for energy. Future investigation into the cellular distribution of lipid droplet accumulation under conditions of dLdh induced changes in aging will be highly illuminating. Overall, our results suggest aberrant lipid metabolism caused by neuronal and glial *dLdh* upregulation may be a contributing factor to decreased lifespan.

### dLdh involvement in long-term memory is age and cell-type dependent

In this study we show that neuronal or glial downregulation or neuronal upregulation of *dLdh* in *D. melanogaster* each cause a deficit in long-term courtship memory but only with age. On the other hand, we found glial *dLdh* upregulation had no impact on long-term courtship memory. We had expected that these interventions would modify glia-neuron lactate shuttling and alter memory in a manner similar to that observed following disruption to the ANLS in vertebrates such as mice, rats, and chicken [39–47,148]. Only one study has examined the impact of the ANLS on cognition with age. That study measured the indirect involvement of the ANLS in memory by knocking out β2-adrenergic receptors in mouse astrocytes to prevent epinephrine signaling induced lactate release [149]. β2-adrenergic receptor astrocyte knockout mice had no change in memory, assessed by the Morris water maze, at a young age, but developed memory deficits at an older age [149]. Interestingly, the memory deficits we uncovered in *dLdh* transgenic flies were similarly restricted to aged animals. These findings suggest that cognitive processing in the aged brain is more susceptible to changes in either the production or shuttling of lactate between glia and neurons compared to the young brain.

In this study we used a fly behavioural paradigm testing cis-vaccenyl acetate (cVA)-retrievable courtship memory, which has neural circuitry that differs from associative courtship memory processing and is likely more complicated than aversive olfactory associative memory [150]. Moreover, we focused on long-term memory because the ANLS has been shown in mice to be dispensable for short-term memory in certain paradigms [40]. Other studies have found manipulating glia causes age-related impairment of aversive olfactory associative memory [53,54]. We found that both glial and neuronal *dLdh* downregulation caused impairment of long-term courtship memory only in aged flies. If glial and neuronal *dLdh* downregulation reduce the ability of glia to release lactate and neurons to oxidize lactate, then these results may reflect the increased dependence on glia-neuron lactate shuttling to fuel long-term courtship memory with age. Perhaps mushroom body neurons encoding long-term courtship memory [85] increase mitochondrial energy flux, as has been proposed for long-term aversive olfactory associative memory encoding [50], but oxidation of lactate becomes a more important mitochondrial fuel supply as flies age. Moreover, an increase in glia-neuron lactate shuttling with age in mushroom bodies would explain why the levels of dLdh increase in the brain with age and why *dLdh* is enriched in mushroom bodies [72]. On the other hand, we also found overexpression of *dLdh* in neurons causes age-related impairment of long-term courtship memory. Neurons with an overabundance of dLdh should retain the ability to oxidize lactate, so a lack of lactate provided by glia seems to be an unlikely explanation for memory deficits in flies with neuronal *dLdh* upregulation. It is possible that excess lactate oxidation by dLdh in neurons promotes increased mitochondrial reactive oxygen species (ROS) production (Young et al, Redox Biology-2020) and potentiates age-related neurodegeneration and memory decline. Interestingly, fumarate was the only metabolite that exhibited an elevation in levels that inversely correlated with memory in aged *dLdh* transgenic flies. The excessive buildup of fumarate in humans with Fumarate hydratase (FH) deficiency, a rare genetic disease, results in brain atrophy, neurologic abnormalities, and intellectual impairment [151,152]. Interestingly, cancer cells with an FH deficiency produce excess fumarate that, in turn, promotes increased ROS levels (Sullivan et al, Mol Cell −2013). Whether mitochondrial lactate oxidation and/or fumarate accumulation associated with altered *dLdh* expression contributes to memory loss or ROS production awaits further investigation. Overall, our findings suggest that *D. melanogaster* long-term courtship memory maintenance with age requires dLdh levels to be tightly regulated in neurons and at or above a critical level in glia.

## Conclusion

In this study we demonstrate the importance of maintaining appropriate levels of dLdh in *D. melanogaster* glia and neurons for maintenance of long-term courtship memory and survival with age. In addition, our results implicate lipid metabolism, 2HG accumulation, and changes in TCA cycle activity as factors underlying the age-related impacts of perturbed *dLdh* expression, which likely modifies glia-neuron lactate shuttling in the fly brain. Moreover, we demonstrate that normal features of *D. melanogaster* aging likely include retention of the ability to form long-term courtship memory, unlike other types of memory, and the accumulation of 2HG in the brain. This study provides a greater understanding of how aging, lactate metabolism, and cognitive processing intersect. Future work is required to determine how lactate metabolism may differ across various *D. melanogaster* brain regions with age and which subtypes of neurons and glia are most susceptible to glia-neuron lactate shuttle perturbations related to accumulation of neutral lipids and the 2HG enantiomers. Our findings highlight a connection between glia-neuron metabolic coupling and age-related memory impairment in *D. melanogaster,* a model highly amenable to further exploration of the role of metabolic coupling on memory processes with age.

## Material and Methods

### Key resources table

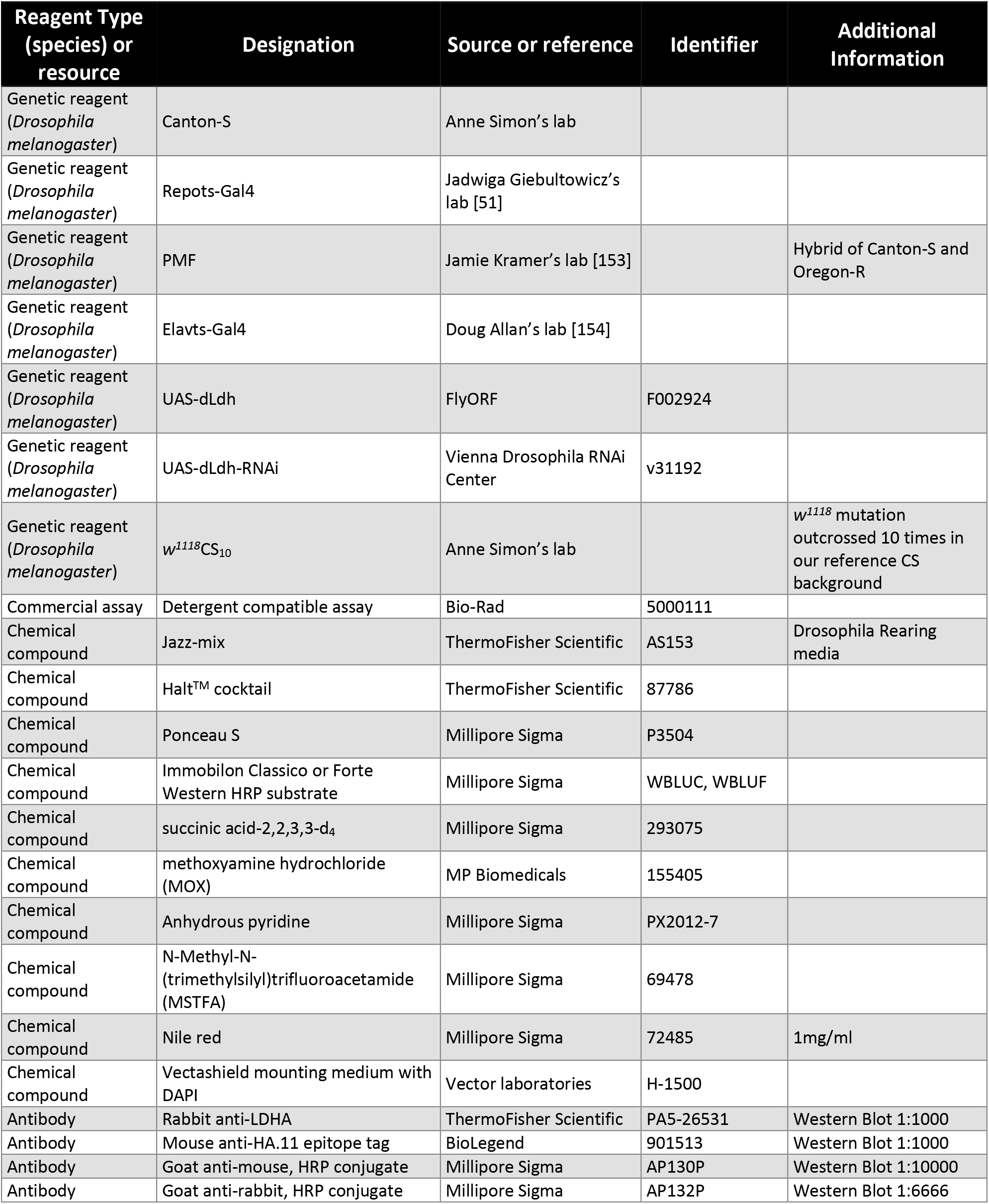
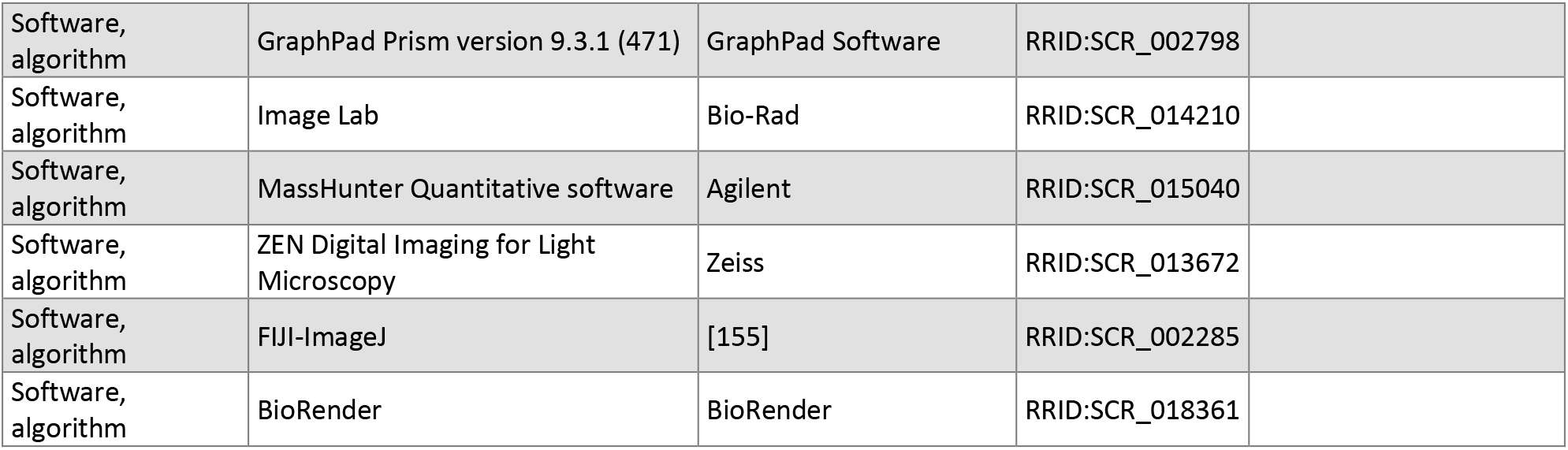

| Reagent Type<br>(species) or<br>resource | Designation | Source or reference | Identifier | Additional<br>Information |
| --- | --- | --- | --- | --- |
| Genetic reagent<br>(*Drosophila melanogaster*) | Canton-S | Anne Simon's lab |  |  |
| Genetic reagent<br>(*Drosophila melanogaster*) | Repots-Gal4 | Jadwiga Giebultowicz's lab [51] |  |  |
| Genetic reagent<br>(*Drosophila melanogaster*) | PMF | Jamie Kramer's lab [153] |  | Hybrid of Canton-S and Oregon-R |
| Genetic reagent<br>(*Drosophila melanogaster*) | Elavts-Gal4 | Doug Allan's lab [154] |  |  |
| Genetic reagent<br>(*Drosophila melanogaster*) | UAS-dLdh | FlyORF | F002924 |  |
| Genetic reagent<br>(*Drosophila melanogaster*) | UAS-dLdh-RNAi | Vienna Drosophila RNAi Center | v31192 |  |
| Genetic reagent<br>(*Drosophila melanogaster*) | *w<sup>1118</sup>*CS<sub>10</sub> | Anne Simon's lab |  | *w<sup>1118</sup>* mutation outcrossed 10 times in our reference CS background |
| Commercial assay | Detergent compatible assay | Bio-Rad | 5000111 |  |
| Chemical compound | Jazz-mix | ThermoFisher Scientific | AS153 | Drosophila Rearing media |
| Chemical compound | Halt™ cocktail | ThermoFisher Scientific | 87786 |  |
| Chemical compound | Ponceau S | Millipore Sigma | P3504 |  |
| Chemical compound | Immobilon Classico or Forte Western HRP substrate | Millipore Sigma | WBLUC, WBLUF |  |
| Chemical compound | succinic acid-2,2,3,3-d<sub>4</sub> | Millipore Sigma | 293075 |  |
| Chemical compound | methoxyamine hydrochloride (MOX) | MP Biomedicals | 155405 |  |
| Chemical compound | Anhydrous pyridine | Millipore Sigma | PX2012-7 |  |
| Chemical compound | N-Methyl-N-(trimethylsilyl)trifluoroacetamide (MSTFA) | Millipore Sigma | 69478 |  |
| Chemical compound | Nile red | Millipore Sigma | 72485 | 1mg/ml |
| Chemical compound | Vectashield mounting medium with DAPI | Vector laboratories | H-1500 |  |
| Antibody | Rabbit anti-LDHA | ThermoFisher Scientific | PA5-26531 | Western Blot 1:1000 |
| Antibody | Mouse anti-HA.11 epitope tag | BioLegend | 901513 | Western Blot 1:1000 |
| Antibody | Goat anti-mouse, HRP conjugate | Millipore Sigma | AP130P | Western Blot 1:10000 |
| Antibody | Goat anti-rabbit, HRP conjugate | Millipore Sigma | AP132P | Western Blot 1:6666 |

|  |  |  |  |
| --- | --- | --- | --- |
| Software, algorithm | GraphPad Prism version 9.3.1 (471) | GraphPad Software | RRID:SCR_002798 |
| Software, algorithm | Image Lab | Bio-Rad | RRID:SCR_014210 |
| Software, algorithm | MassHunter Quantitative software | Agilent | RRID:SCR_015040 |
| Software, algorithm | ZEN Digital Imaging for Light Microscopy | Zeiss | RRID:SCR_013672 |
| Software, algorithm | FIJI-ImageJ | [155] | RRID:SCR_002285 |
| Software, algorithm | BioRender | BioRender | RRID:SCR_018361 |

### Fly stocks and maintenance

*Drosophila melanogaster* fly stocks were maintained on a 12:12-h light:dark cycle at 50% relative humidity on Jazz-mix (AS153, ThermoFisher Scientific) food or food made in-house using an equivalent recipe. All experiments were done using male flies exclusively. Flies undergoing negative geotaxis assays were maintained in vials (AS507, ThermoFisher Scientific). Flies for all other experiments were maintained under individual housing conditions, each within a 2ml deep well of a 96 well plate (89237-526, VWR) half filled with food. Flies in 96 well plates were transferred to new food at least once every two weeks. In house Canton-S flies were used as non-transgenic controls, maintained at 25°C for courtship conditioning and 25°C or 29°C for survival analysis. All transgenic flies were raised at 18°C in order to prevent activation of the temporal and regional gene expression targeting (TARGET) system then maintained at 29°C post-eclosion for experimental purposes [89]. The fly stocks used in this study were as follows: Canton-S (CS), UAS-dLdh (F002924; FlyORF), UAS-dLdh-RNAi (v31192; VDRC), Elavts-Gal4 (w,elavC155-Gal4,UAS-Dicer2;tuGal80ts,UAS-nGFP; Provided by Doug Allan, University of British Columbia), Repots-Gal4 (w; tubGal80TS;Repo-Gal4; Provided by Jaga Giebultowicz, Oregon State University), PMF (Hybrid of Canton-S and Oregon-R; Provided by Jamie Kramer, Dalhousie University), and w^1118^_CS10_. UAS-dLdh and UAS-dLdh-RNAi were outcrossed for five generations to w^1118^_CS10_, a *w*^1118^ mutation outcrossed 10 times in our reference CS background.

### Survival analysis

Adult CS male flies were collected with or without brief CO2 anesthesia and maintained in 96 well plates as described above. Those flies were raised at 25°C, and their survival assayed at 29°C or 25°C in three separate trials. For each temperature group, a survival curve was generated by consolidating data from all three trials on that group. Transgenic flies and their associated control groups lacking UAS constructs were each assessed in a trial of groups tested at 29°C, after being raised at 18°C, and all collected within a week of hatching from pupae. For all flies, the first day in 29°C conditions was considered the first day of survival. Deaths were recorded 3-4 times per week. Flies which managed to escape were censored. Catastrophic death events where greater than 20% of flies suddenly die at a rate unrepresentative of the rest of the population or among repeated independent trials were excluded [156]. Survival curves were generated using percentage survival computed with the product limit (Kaplan-Meier) method for each group of flies and censored flies omitted from the graph. To compare survival curves, the log-rank (Mantel-Cox) test was used to determine differences in curves overall, median survival, maximum lifespan, and rate of dying (hazard ratio with 95% confidence interval calculated using the Mantel-Haenszel method). All calculations were done using GraphPad Prism version 9.3.1 (RRID:SCR_002798).

### Western blot

Frozen flies were decapitated and heads in a 2% sodium dodecyl sulfate (SDS) lysis buffer made with 50mM Tris, 1mM Ethylenediaminetetraacetic acid (EDTA) and protease inhibitors, 1mM sodium orthovanadate, 1mM phenylmethylsulfonyl fluoride, and Halt™ cocktail (87786, ThermoFisher Scientific). After homogenization with a pestle and sonication of the heads, the samples were centrifuged and protein in the supernatant was extracted. The concentration of extracted protein was determined using a detergent compatible assay (5000111, Bio-Rad). Extracted protein was combined with a bromophenol blue based loading buffer, boiled, resolved on an acrylamide gel and transferred to a polyvinylidene fluoride (PVDF) membrane. Total protein level was measured by staining with 0.1%

Ponceau S (P3504, Millipore Sigma) in 5% glacial acetic acid for 3 minutes with excess Ponceau S removed with a 3-minute methanol wash prior to imaging. Blots were then destained, using 5-minute washes with tris buffered saline containing tween 20 (TBST) four times, and blocked with bovine serum albumin (BSA) and milk in TBST for at least 1 hour. Blots were probed with primary antibody at 4°C overnight. Primary antibodies were as follows: rabbit anti-LDHA (PA5-26531, ThermoFisher Scientific; 1:1000), and mouse anti-HA.11 epitope tag (901513, BioLegend; 1:1000). Subsequently, blots were probed with horse radish peroxidase (HRP) conjugated secondary antibodies at room temperature for 2 hours. Secondary antibodies were as follows: goat anti-mouse (AP130P, Millipore Sigma; 1:10000), and goat anti-rabbit (AP132P, Millipore Sigma; 1:6666). Antibodies were made in TBST with 0.01% sodium azide, and blots were washed in TBST before and after each probe. Immobilon Classico or Forte Western HRP substrate (WBLUC, WBLUF, Millipore Sigma) were used to detect chemiluminescent signals on probed blots. Chemiluminescence and Ponceaus S signal density were each imaged using a ChemiDoc XRS imaging system (170-8070, Bio-Rad) and quantified using Image Lab software (RRID:SCR_014210). For quantification purposes, western blot band intensity was standardized to total Ponceaus S signal density for each lane.

### Metabolite Analysis using Gas Chromatography-Mass Spectrometry (GC-MS)

Frozen fly heads were stored at −80°C. Prior to analysis, heads were transferred to pre-tared 2 mL screw cap tubes containing 1.4 mm ceramic beads (15340153, ThermoFisher Scientific), the mass was recorded, and samples were placed in liquid nitrogen. As previously described [157], a solution of 90% methanol (34860, Millipore Sigma) containing a succinic acid-2,2,3,3-d_4_ (293075, Millipore Sigma) standard was added to each sample and homogenized at 4°C for 30 seconds using an Omni BeadRuptor 24 set at 6.45 m/s. The samples were incubated at −20°C for an hour, centrifuged at 10,000xg at 4°C, and the supernatant was transferred to a 1.5 ml microfuge tube. The sample was dried overnight using a Thermo Savant vacuum centrifuge (SPD130DLX-115) set to ambient temperature and attached a Savant Refrigerated Vapor Trap (RVT5105-230). Samples were resuspended and derivatized by adding 40 μl of 40 mg/ml methoxyamine hydrochloride (MOX; 155405, MP Biomedicals) in anhydrous pyridine (PX2012-7, Millipore Sigma) and incubated at 35°C for one hour with shaking at 600 rpm using an Eppendorf ThermoMixer F1.5. Then 25 μl of the supernatant was transferred into a 2 ml sample vial containing a 250 μl deactivated glass insert (5181-8872, Agilent). 40 μl of N-Methyl-N- (trimethylsilyl)trifluoroacetamide (MSTFA; 69478, Millipore Sigma) was added and the vial placed in a Benchmark Multi-Therm heat shaker set at 40°C and 250 rpm. Following the incubation period, samples were immediately analyzed using a 7890B-5977B MSD Agilent GC-MS. 1 μl of the derivatized sample was injected into a 0.25 mm i.d., 30-meter DB-5MS column (Agilent) and the split ratio was set to 50:1. Initial oven temperature was set to 95°C with one minute of hold time. The first ramp was set to 40°C/min until 110°C, then 5°C/min until it reaches 250°C and finally 25°C/min until 330°C. The area under the peak for each metabolite was calculated using MassHunter Quantitative software (RRID: SCR_015040). The value for each metabolite was normalized to the succinic acid-2,2,3,3-d4 internal standard and sample mass. For each condition, outliers were identified and removed using the Robust regression and Outlier removal (ROUT) method [158]. Relative abundance of each metabolite was calculated by further normalizing to the average for control flies at 7 days of age.

### Courtship conditioning

In order to prevent male flies from having any courtship experience prior to training, they were isolated in 96 well plates either early post-eclosion, and briefly anesthetized on ice, or as pupae just prior to eclosion. Long-term courtship memory is formed by males that learn to reduce courtship to a target pre-mated female (PMF) by a training period in which the male experiences constant rejection by a PMF. PMF or CS females, aged five days in vials at 25°C in the presence of males, were collected with CO2 anesthesia one to five hours prior to either training or testing in order to act as the target pre-mated female. Males were trained by transferring them to a new well of a 96 well plate with a PMF for seven to eight hours. Naïve males were sham trained by transferring to a new well of a 96 well plate alone. 20-24 hours after training, long-term courtship memory testing was performed by providing a target PMF to each male in a transparent circular enclosure for 10 minutes without food. Training and testing were done under the same temperature and humidity conditions males were aged under. Video recordings of courtship during the 10 minutes testing of both naïve and trained males were generated using an iPad (A1458, Apple) or iPad mini (A1489, Apple) and scored manually, with the scorer blind to training status, genotype, and age. Male behaviours considered courtship include orienting, chasing, tapping, licking, vibrating wings, unilateral wing extension, and abdomen bending in attempt to copulate. Courtship index (CI) was calculated for each tested male as the percent of time exhibiting courtship behaviour. When CI of trained males is lower than CI of naïve males within the same age and genotype this is considered an exhibition of long-term courtship memory. Memory Indexes (MI) were calculated using CI for each trained fly compared to the average of naïve flies in the same condition [MI = 1 – (CI_Trained_/CI_Average Naive_)].

### Negative geotaxis climbing ability assays

Male flies were raised in vials in groups of 50 with surviving flies tested weekly. Flies were transferred to new vials of food after each test to be retested the following week, up until 28 days of age. Climbing ability was assayed at 25°C and 50% humidity in a counter-current apparatus according to an adapted protocol [159,160]. For each test, flies were transferred to a vertically oriented plastic conical vial (352017, Corning) on the bottom of the counter current device. Flies were then tapped down 6-7 times to bring them down to the bottom of the vial. With all flies on the bottom of the vial, the upper portion of the counter-current device was displaced to open access to a clean empty plastic conical vial on the top of the apparatus. Flies will climb upwards due to their natural inclination for negative geotaxis. After 10 seconds of letting flies climb upwards towards the top vial, the upper portion of the counter-current device was displaced again to separate flies that had reached the top vial from flies that had not. A performance index (PI) was calculated by dividing the number of flies which had reached the top vial within 10 seconds by the total number of flies originally placed in the bottom vial [PI = # Top Flies / Total # Flies].

### Nile red staining

Brains of male flies aged 21 days at 29°C were analyzed. Brains were dissected and stained with Nile Red for neutral lipids according to a protocol adapted from [91]. Flies were briefly anesthetized with carbon dioxide and brains dissected on ice cold phosphate buffered saline (PBS). Brains were fixed in 4% paraformaldehyde and 0.5% Triton X in PBS for 45 minutes on ice, then washed three times in PBS on ice for 15 minutes. Brains were stained with 1μg/ml rile red in PBS on ice, freshly made from a 1mg/ml stock of nile red (72485, Millipore Sigma) dissolved in acetone stock solution. After staining, brains were rinsed once in PBS on ice, mounted onto glass slides using Vectashield mounting medium with DAPI (H-1500, Vector Laboratories), and imaged by confocal microscopy on the same day.

### Confocal Microscopy

For Nile Red staining, fly brains were imaged on an inverted Zeiss LSM 800 scanning laser confocal microscope with an LCI Plan-Neofluar 25x/0.8 Imm Korr DIC water immersion objective (Zeiss). Tiles of Z-stacks with 10μm intervals were recorded spanning the entire brain. Nile Red and DAPI fluorescence were sequentially imaged. Nile Red fluorescence was detected using 559μm excitation and 646μm emission acquired with a 53μm pinhole. DAPI fluorescence was detected using 353μm excitation and 465μm emission acquired with a 52μm pinhole. Maximum intensity projections were generated for analysis and visualization (RRID:SCR_013672, ZEN Digital Imaging for Light Microscopy). Nile red mean fluorescence in the total area of each brain and in a square sample of the background was quantified in FIJI-ImageJ (Schindelin et al., 2012; RRID:SCR_002285) using the protocol described by [161]. Background was subtracted from brain signal for each brain and normalized to the average of control brains.

### Statistical analyses

Data are presented as mean ± SEM unless otherwise specified and were analyzed statistically and visualized using GraphPad Prism version 9.3.1 (471) (RRID:SCR_002798). All raw data and statistical analyses are available on the Dryad data repository (https://doi.org/10.5061/dryad.2v6wwpzsb).

## Acknowledgements

We thank the Biotron Facility, Insect suite at Western University, where the flies were reared in specialized climate-controlled walk-in rooms (25°C), or in temperature and humidity-controlled incubators (18°C or 29°C). We thank Amina Kassam, Delina Narendran, Sanjana Arora, Ikjot Mann, Jenan Altoum, Kevin Lee, and Sanjit Sumal for assisting in manual scoring of male fly courtship. We thank Jamie Kramer for assistance in planning the courtship conditioning paradigm and lending us the enclosures to video courtship in.

## Author Details

Funding support for AKF and RCC was provided by the Natural Sciences and Engineering Research Council of Canada under award number RGPIN-2019-355803. RCC was supported by the Helen Battle Professorship. AKF received an Ontario Graduate Scholarship and Postgraduate Scholarship – Doctoral from the Natural Sciences and Engineering Research Council of Canada.

Support for JWR and AFS was provided by the Natural Sciences and Engineering Research Council of Canada under award number RGPIN-2015-04275 and RGPIN-2022-05054. JWR received an Ontario Graduate Scholarship and Postgraduate Scholarship – Doctoral from the Natural Sciences and Engineering Research Council of Canada.

Support for NHM and JMT was provided by the National Institute of General Medical Sciences of the National Institutes of Health under award number R35GM119557.

---

**Figure S1.**
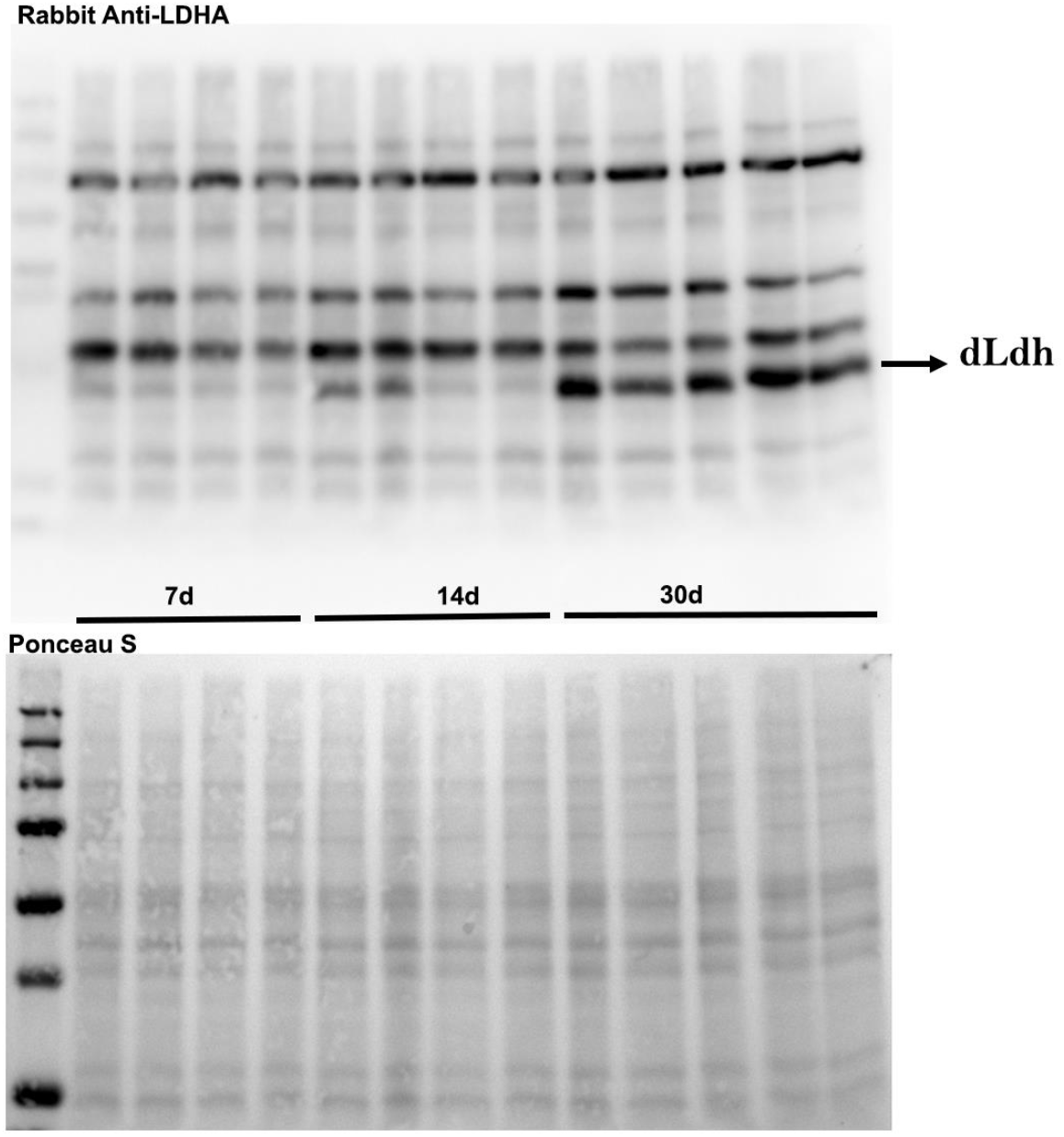
Full western blot images for Figure 1E dLdh detection. Full images for western blot analysis to detect dLdh (~35.5kD) in protein extracts from heads of male Canton-S flies aged 7, 14 or 30 days at 29°C quantified in Figure 1E. Ponceau S stained blot revealing all transferred proteins is shown below.

**Figure S2.**
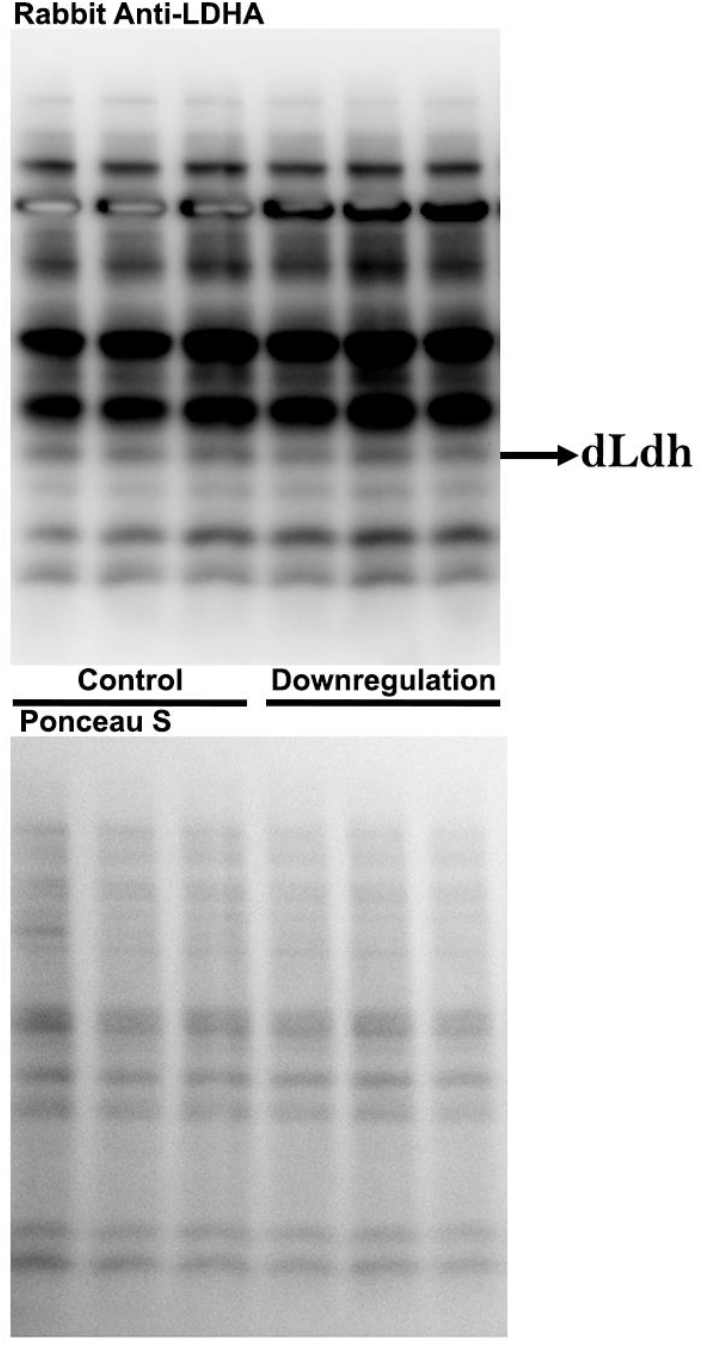
Full western blot images for Figure 2A dLdh detection. Full images for western blot analysis to detect dLdh (~35.5kD) in head extracts from neuronal transgenic male flies aged 21 days at 29°C quantified in Figure 2A. Ponceau-S stained blot revealing all transferred proteins is shown below.

**Figure 3S.**
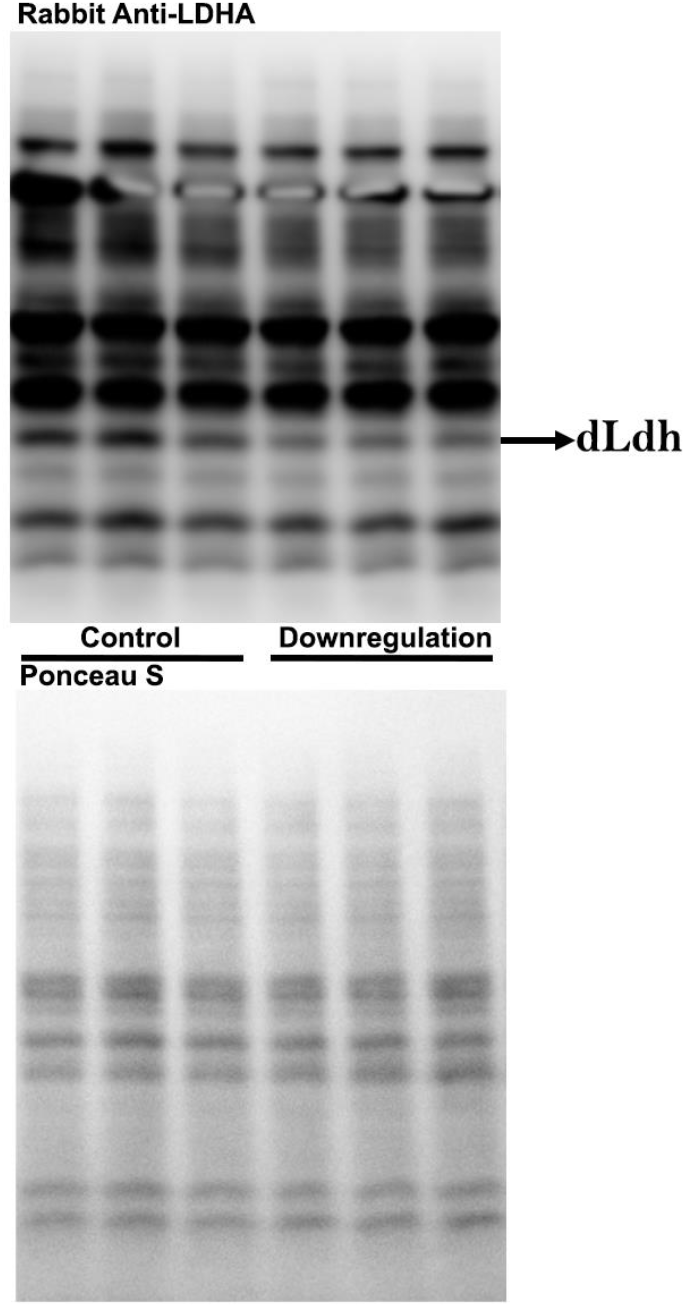
Full western blot images for Figure 3A dLdh detection. Full images for western blot analysis to detect dLdh (~35.5kD) in head extracts from glial transgenic male flies aged 21 days at 29°C quantified in Figure 3A. Ponceau-S stained blot revealing all transferred proteins is shown below.

**Figure 4S.**
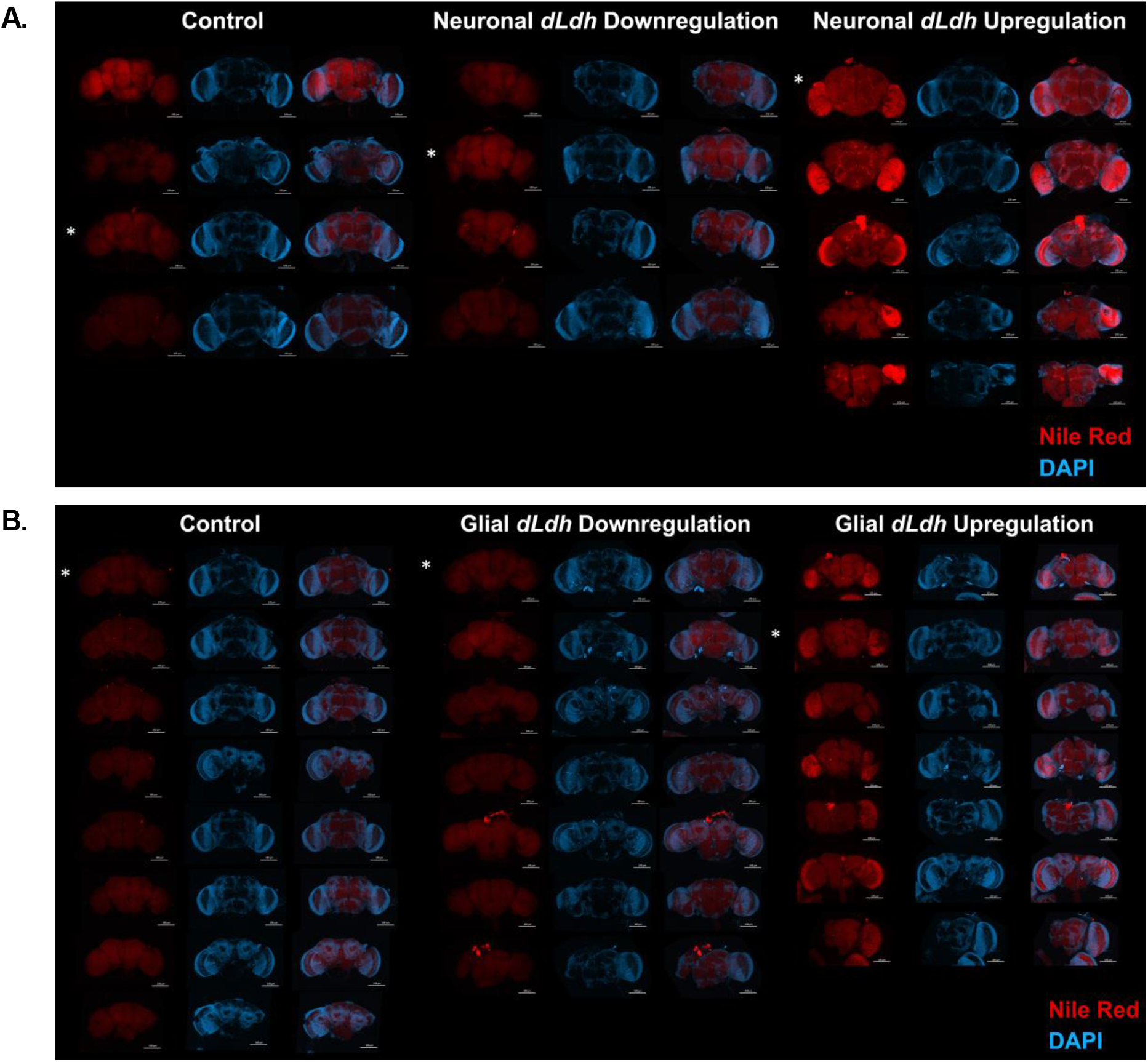
Neutral lipid staining of all 21 day aged male transgenic fly whole brains with altered glial or neuronal *dLdh* expression. **A.** All confocal fluorescence microscopy images of neuronal *dLdh* transgenic fly brains removed at 21 days of age at 29°C and stained for DNA (Blue = DAPI) and neutral lipids (Red = Nile Red). **B.** All confocal fluorescence microscopy images of glial *dLdh* transgenic fly brains removed at 21 days of age at 29°C and stained for DNA (Blue = DAPI) and neutral lipids (Red = Nile Red).

**Figure 5S.**
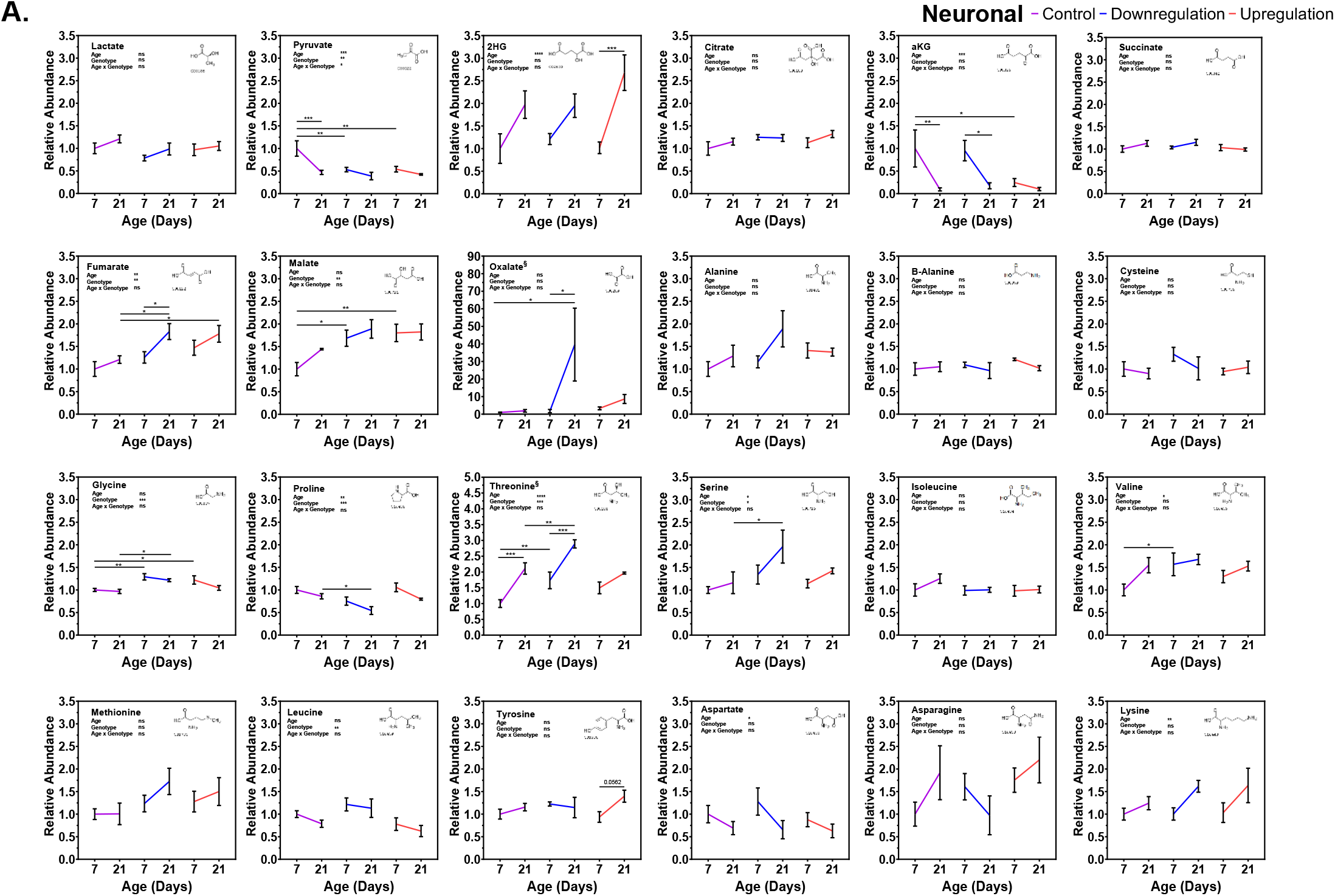
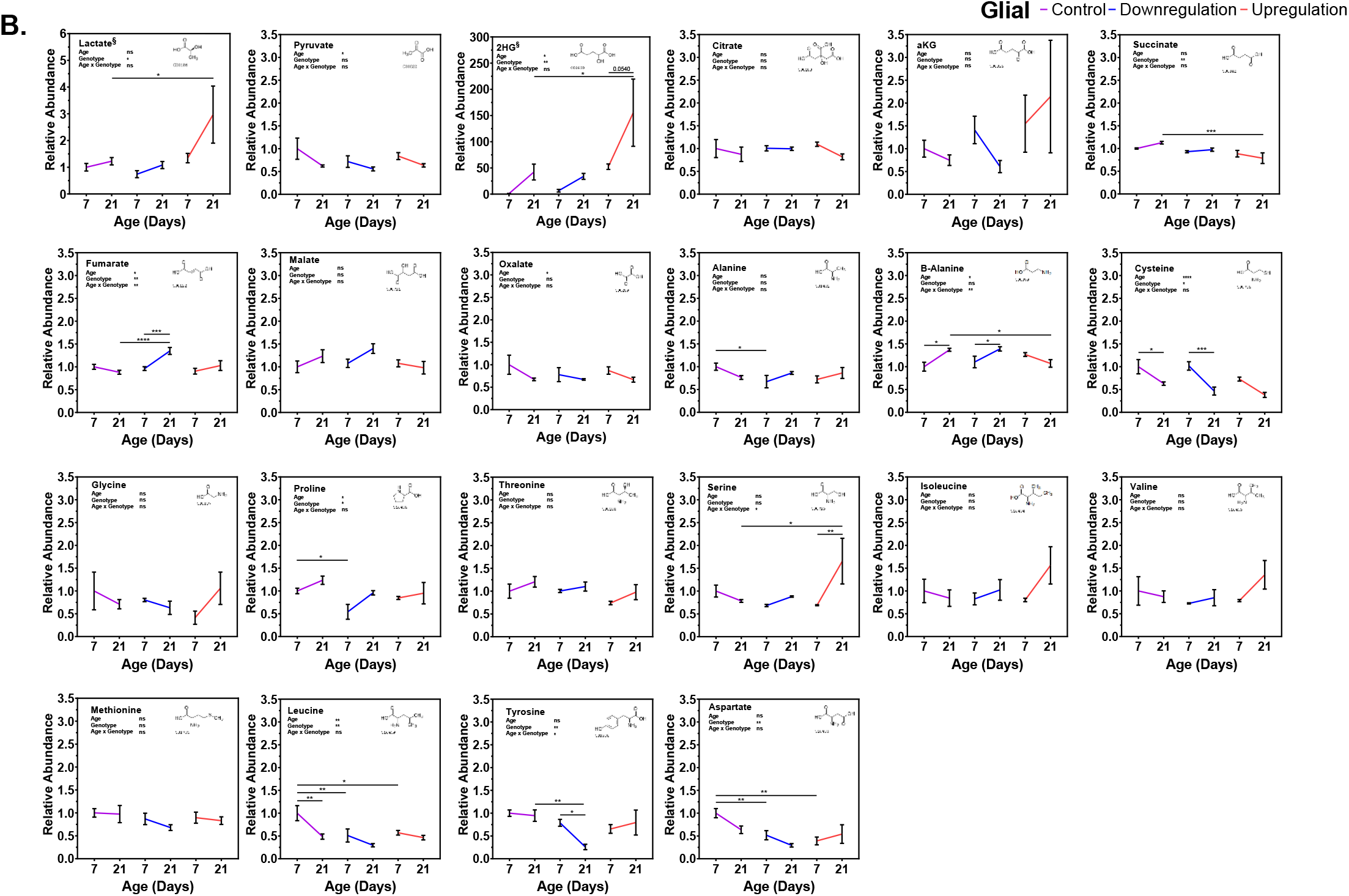
Expanded panel of metabolites detected in transgenic male fly heads aged 7 and 21 days with altered expression of neuronal or glial *dLdh*. **A.** All metabolites measured by gas chromatography-mass spectrometry (GC-MS) in the heads of neuronal *dLdh* transgenic male flies aged 7 and 21 days at 29° **B.** All metabolites measured by gas chromatography-mass spectrometry (GC-MS) in the heads of glial *dLdh* transgenic male flies aged 7 and 21 days at 29°. Comparisons for each metabolite were done between genotype and age groups of neuronal and glial *dLdh* transgenic flies separately using two-way ANOVAs with Dunnet’s multiple comparisons with control for each age group and with Šídák’s multiple comparisons between age groups within each genotype. Effects of age, genotype, and age by genotype interaction are denoted on the top left of each graph. The structural formula for each metabolite were obtained from the Kyoto Encyclopedia of Genes and Genomes (KEGG) chemical compound database with associated C number below.

**Figure 6S.**
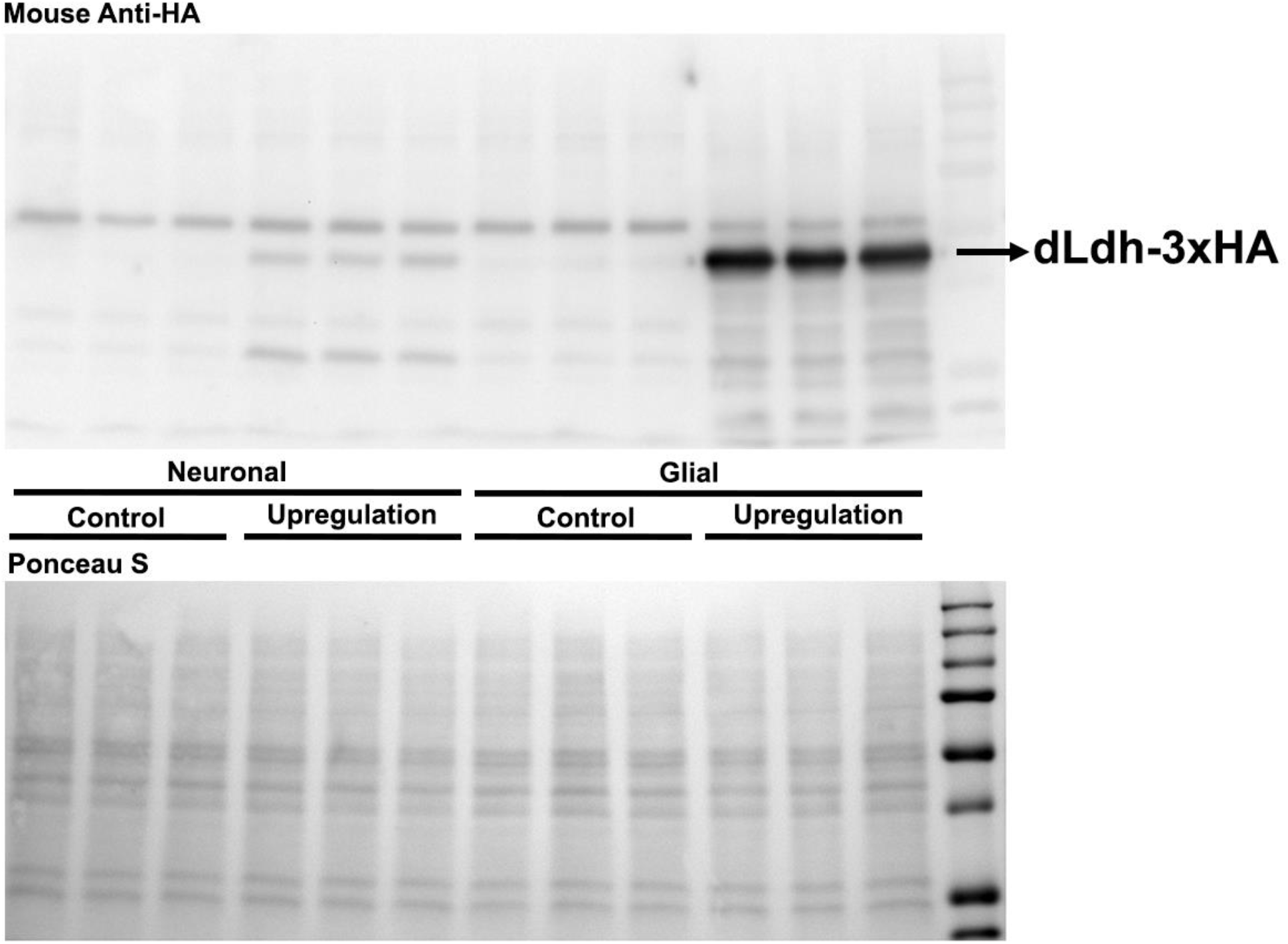
Full western blot images for Figure 2A and 3A dLdh-3xHA detection.

## References

[1] Mery F, Kawecki TJ. A Cost of Long-Term Memory in Drosophila. Science (80-) 2005;308:1148–1148. PMID:15905396. DOI:10.1126/science.1111331.

[2] Laughlin SB, de Ruyter van Steveninck RR, Anderson JC. The metabolic cost of neural information. Nat Neurosci 1998;1:36–41. DOI:10.1038/236.

[3] Mink JW, Blumenschine RJ, Adams DB. Ratio of central nervous system to body metabolism in vertebrates: Its constancy and functional basis. Am J Physiol - Regul Integr Comp Physiol 1981;10:203–12. PMID:7282965. DOI:10.1152/ajpregu.1981.241.3.r203.

[4] Harris JJ, Jolivet R, Attwell D. Synaptic Energy Use and Supply. Neuron 2012;75:762–77. PMID:22958818. DOI:10.1016/j.neuron.2012.08.019.

[5] Brooks GA, Arevalo JA, Osmond AD, Leija RG, Curl CC, Tovar AP. Lactate in contemporary biology: a phoenix risen. J Physiol 2021;35:JP280955. DOI:10.1113/JP280955.

[6] Ivanov A, Mukhtarov M, Bregestovski P, Zilberter Y. Lactate Effectively Covers Energy Demands during Neuronal Network Activity in Neonatal Hippocampal Slices. Front Neuroenergetics 2011;3:1–14. DOI:10.3389/fnene.2011.00002.

[7] Bouzier-Sore A-K, Voisin P, Canioni P, Magistretti PJ, Pellerin L. Lactate is a Preferential Oxidative Energy Substrate over Glucose for Neurons in Culture. J Cereb Blood Flow Metab 2003;23:1298–306. PMID:14600437. DOI:10.1097/01.WCB.0000091761.61714.25.

[8] Wyss MT, Jolivet R, Buck A, Magistretti PJ, Weber B. In Vivo Evidence for Lactate as a Neuronal Energy Source. J Neurosci 2011;31:7477–85. PMID:21593331. DOI:10.1523/JNEUROSCI.0415-11.2011.

[9] Poitry-Yamate CL, Poitry S, Tsacopoulos M. Lactate released by Muller glial cells is metabolized by photoreceptors from mammalian retina. J Neurosci 1995;15:5179–91. PMID:7623144. DOI:10.1523/jneurosci.15-07-05179.1995.

[10] Schurr A, West CA, Rigor BM. Lactate-Supported Synaptic Function in the Rat Hippocampal Slice Preparation. Science (80-) 1988;240:1326–8. PMID:3375817. DOI:10.1126/science.3375817.

[11] Larrabee MG. Lactate metabolism and its effects on glucose metabolism in an excised neural tissue. J Neurochem 1995;64:1734–41. PMID:7891102. DOI:10.1046/j.1471-4159.1995.64041734.x.

[12] Karagiannis A, Gallopin T, Lacroix A, Plaisier F, Piquet J, Geoffroy H, et al. Lactate is an energy substrate for rodent cortical neurons and enhances their firing activity. Elife 2021;10:1–40. PMID:34766906. DOI:10.7554/eLife.71424.

[13] Fink K, Velebit J, Vardjan N, Zorec R, Kreft M. Noradrenaline-induced l-lactate production requires d-glucose entry and transit through the glycogen shunt in single-cultured rat astrocytes. J Neurosci Res 2021:jnr.24783. DOI:10.1002/jnr.24783.

[14] Sorg O, Magistretti PJ. Characterization of the glycogenolysis elicited by vasoactive intestinal peptide, noradrenaline and adenosine in primary cultures of mouse cerebral cortical astrocytes. Brain Res 1991;563:227–33. PMID:1664773. DOI:10.1016/0006-8993(91)91538-C.

[15] Mächler P, Wyss MT, Elsayed M, Stobart J, Gutierrez R, von Faber-Castell A, et al. In Vivo Evidence for a Lactate Gradient from Astrocytes to Neurons. Cell Metab 2016;23:94–102. PMID:26698914. DOI:10.1016/j.cmet.2015.10.010.

[16] Ruminot I, Schmälzle J, Leyton B, Barros LF, Deitmer JW. Tight coupling of astrocyte energy metabolism to synaptic activity revealed by genetically encoded FRET nanosensors in hippocampal tissue. J Cereb Blood Flow Metab 2019;39:513–23. PMID:29083247. DOI:10.1177/0271678X17737012.

[17] Zuend M, Saab AS, Wyss MT, Ferrari KD, Hösli L, Looser ZJ, et al. Arousal-induced cortical activity triggers lactate release from astrocytes. Nat Metab 2020;2:179–91. DOI:10.1038/s42255-020-0170-4.

[18] Pellerin L, Magistretti PJ. Sweet sixteen for ANLS. J Cereb Blood Flow Metab 2012;32:1152–66. PMID:22027938. DOI:10.1038/jcbfm.2011.149.

[19] Pellerin L, Magistretti PJ. Glutamate uptake into astrocytes stimulates aerobic glycolysis: a mechanism coupling neuronal activity to glucose utilization. Proc Natl Acad Sci U S A 1994;91:10625–9. PMID:7938003..

[20] Dienel GA. Brain lactate metabolism: the discoveries and the controversies. J Cereb Blood Flow Metab 2012;32:1107–38. PMID:22186669. DOI:10.1038/jcbfm.2011.175.

[21] Dienel GA. Lack of appropriate stoichiometry: Strong evidence against an energetically important astrocyte-neuron lactate shuttle in brain. J Neurosci Res 2017;95:2103–25. PMID:28151548. DOI:10.1002/jnr.24015.

[22] Drulis-Fajdasz D, Gizak A, Wójtowicz T, Wiśniewski JR, Rakus D. Aging-associated changes in hippocampal glycogen metabolism in mice. Evidence for and against astrocyte-to-neuron lactate shuttle. Glia 2018;66:1481–95. PMID:29493012. DOI:10.1002/glia.23319.

[23] Bak LK, Walls AB. CrossTalk opposing view: lack of evidence supporting an astrocyte-to-neuron lactate shuttle coupling neuronal activity to glucose utilisation in the brain. J Physiol 2018;596:351–3. PMID:29292507. DOI:10.1113/JP274945.

[24] Chih C-P, Roberts EL. Energy Substrates for Neurons during Neural Activity: A Critical Review of the Astrocyte-Neuron Lactate Shuttle Hypothesis. J Cereb Blood Flow Metab 2003;23:1263–81. PMID:14600433. DOI:10.1097/01.WCB.0000081369.51727.6F.

[25] Díaz-García CM, Mongeon R, Lahmann C, Koveal D, Zucker H, Yellen G. Neuronal Stimulation Triggers Neuronal Glycolysis and Not Lactate Uptake. Cell Metab 2017;26:361–374.e4. PMID:28768175. DOI:10.1016/j.cmet.2017.06.021.

[26] Lundgaard I, Li B, Xie L, Kang H, Sanggaard S, Haswell JDR, et al. Direct neuronal glucose uptake heralds activity-dependent increases in cerebral metabolism. Nat Commun 2015;6:6807. PMID:25904018. DOI:10.1038/ncomms7807.

[27] Ng FS, Sengupta S, Huang Y, Yu AM, You S, Roberts MA, et al. TRAP-seq Profiling and RNAi-Based Genetic Screens Identify Conserved Glial Genes Required for Adult Drosophila Behavior. Front Mol Neurosci 2016;9. DOI:10.3389/fnmol.2016.00146.

[28] Freeman MR. Drosophila Central Nervous System Glia. Cold Spring Harb Perspect Biol 2015;7:a020552. PMID:25722465. DOI:10.1101/cshperspect.a020552.

[29] Bittern J, Pogodalla N, Ohm H, Brüser L, Kottmeier R, Schirmeier S, et al. Neuron–glia interaction in the Drosophila nervous system. Dev Neurobiol 2020:dneu.22737. DOI:10.1002/dneu.22737.

[30] Yildirim K, Petri J, Kottmeier R, Klämbt C. Drosophila glia: Few cell types and many conserved functions. Glia 2019;67:5–26. DOI:10.1002/glia.23459.

[31] Tsacopoulos M, Evêquoz-Mercier V, Perrottet P, Buchner E. Honeybee retinal glial cells transform glucose and supply the neurons with metabolic substrate. Proc Natl Acad Sci U S A 1988;85:8727–31. PMID:3186756. DOI:10.1073/pnas.85.22.8727.

[32] Tsacopoulos M, Veuthey A, Saravelos S, Perrottet P, Tsoupras G. Glial cells transform glucose to alanine, which fuels the neurons in the honeybee retina. J Neurosci 1994;14:1339–51. PMID:8120629. DOI:10.1523/JNEUROSCI.14-03-01339.1994.

[33] Tsacopoulos M, Coles JA, Van de Werve G. The supply of metabolic substrate from glia to photoreceptors in the retina of the honeybee drone. J Physiol (Paris) 1987;82:279–87. PMID:3503929..

[34] Volkenhoff A, Weiler A, Letzel M, Stehling M, Klämbt C, Schirmeier S. Glial Glycolysis Is Essential for Neuronal Survival in Drosophila. Cell Metab 2015;22:437–47. PMID:26235423. DOI:10.1016/j.cmet.2015.07.006.

[35] Delgado MG, Oliva C, López E, Ibacache A, Galaz A, Delgado R, et al. Chaski, a novel Drosophila lactate/pyruvate transporter required in glia cells for survival under nutritional stress. Sci Rep 2018;8:1186. PMID:29352169. DOI:10.1038/s41598-018-19595-5.

[36] Liu L, MacKenzie KR, Putluri N, Maletić-Savatić M, Bellen HJ. The Glia-Neuron Lactate Shuttle and Elevated ROS Promote Lipid Synthesis in Neurons and Lipid Droplet Accumulation in Glia via APOE/D. Cell Metab 2017;26:719–737.e6. PMID:28965825. DOI:10.1016/j.cmet.2017.08.024.

[37] González-Gutiérrez A, Ibacache A, Esparza A, Barros LF, Sierralta J. Neuronal lactate levels depend on glia-derived lactate during high brain activity in Drosophila. Glia 2020;68:1213–27. PMID:31876077. DOI:10.1002/glia.23772.

[38] Li F, Sami A, Noristani HN, Slattery K, Qiu J, Groves T, et al. Glial Metabolic Rewiring Promotes Axon Regeneration and Functional Recovery in the Central Nervous System. Cell Metab 2020:1–19. PMID:32941799. DOI:10.1016/j.cmet.2020.08.015.

[39] Gibbs ME, Anderson DG, Hertz L. Inhibition of glycogenolysis in astrocytes interrupts memory consolidation in young chickens. Glia 2006;54:214–22. PMID:17674369. DOI:10.1002/glia.20377.

[40] Suzuki A, Stern SA, Bozdagi O, Huntley GW, Walker RH, Magistretti PJ, et al. Astrocyte-Neuron Lactate Transport Is Required for Long-Term Memory Formation. Cell 2011;144:810–23. PMID:21376239. DOI:10.1016/j.cell.2011.02.018.

[41] Newman LA, Korol DL, Gold PE. Lactate Produced by Glycogenolysis in Astrocytes Regulates Memory Processing. PLoS One 2011;6:e28427. PMID:22180782. DOI:10.1371/journal.pone.0028427.

[42] Boury-Jamot B, Carrard A, Martin JL, Halfon O, Magistretti PJ, Boutrel B. Disrupting astrocyte-neuron lactate transfer persistently reduces conditioned responses to cocaine. Mol Psychiatry 2016;21:1070–6. PMID:26503760. DOI:10.1038/mp.2015.157.

[43] Zhang Y, Xue Y, Meng S, Luo Y, Liang J, Li J, et al. Inhibition of Lactate Transport Erases Drug Memory and Prevents Drug Relapse. Biol Psychiatry 2016;79:928–39. PMID:26293178. DOI:10.1016/j.biopsych.2015.07.007.

[44] Tadi M, Allaman I, Lengacher S, Grenningloh G, Magistretti PJ. Learning-Induced Gene Expression in the Hippocampus Reveals a Role of Neuron - Astrocyte Metabolic Coupling in Long Term Memory. PLoS One 2015;10:e0141568. PMID:26513352. DOI:10.1371/journal.pone.0141568.

[45] Vezzoli E, Calì C, De Roo M, Ponzoni L, Sogne E, Gagnon N, et al. Ultrastructural Evidence for a Role of Astrocytes and Glycogen-Derived Lactate in Learning-Dependent Synaptic Stabilization. Cereb Cortex 2020;30:2114–27. PMID:31807747. DOI:10.1093/cercor/bhz226.

[46] Descalzi G, Gao V, Steinman MQ, Suzuki A, Alberini CM. Lactate from astrocytes fuels learning-induced mRNA translation in excitatory and inhibitory neurons. Commun Biol 2019;2:247. PMID:31925050. DOI:10.1038/s42003-019-0495-2.

[47] Korol DL, Gardner RS, Tunur T, Gold PE. Involvement of lactate transport in two object recognition tasks that require either the hippocampus or striatum. Behav Neurosci 2019;133:176–87. PMID:30907617. DOI:10.1037/bne0000304.

[48] de Tredern E, Rabah Y, Pasquer L, Minatchy J, Plaçais P-Y, Preat T. Glial glucose fuels the neuronal pentose phosphate pathway for long-term memory. Cell Rep 2021;36:109620. DOI:10.1016/j.celrep.2021.109620.

[49] Silva B, Mantha OL, Schor J, Pascual A, Plaçais P-Y, Pavlowsky A, et al. Glia fuel neurons with locally synthesized ketone bodies to sustain memory under starvation. Nat Metab 2022. DOI:10.1038/s42255-022-00528-6.

[50] Plaçais P-Y, de Tredern É, Scheunemann L, Trannoy S, Goguel V, Han K-A, et al. Upregulated energy metabolism in the Drosophila mushroom body is the trigger for long-term memory. Nat Commun 2017;8:15510. PMID:28580949. DOI:10.1038/ncomms15510.

[51] Long DM, Frame AK, Reardon PN, Cumming RC, Hendrix DA, Kretzschmar D, et al. Lactate dehydrogenase expression modulates longevity and neurodegeneration in *Drosophila melanogaster*. Aging (Albany NY) 2020;12:10041–58. DOI:10.18632/aging.103373.

[52] Niccoli T, Kerr F, Snoeren I, Fabian D, Aleyakpo B, Ivanov D, et al. Activating transcription factor 4-dependent lactate dehydrogenase activation as a protective response to amyloid beta toxicity. Brain Commun 2021;3:1–13. DOI:10.1093/braincomms/fcab053.

[53] Matsuno M, Horiuchi J, Ofusa K, Masuda T, Saitoe M. Inhibiting Glutamate Activity during Consolidation Suppresses Age-Related Long-Term Memory Impairment in Drosophila. IScience 2019;15:55–65. DOI:10.1016/j.isci.2019.04.014.

[54] Yamazaki D, Horiuchi J, Ueno K, Ueno T, Saeki S, Matsuno M, et al. Glial Dysfunction Causes Age-Related Memory Impairment in Drosophila. Neuron 2014;84:753–63. PMID:25447741. DOI:10.1016/j.neuron.2014.09.039.

[55] Chinopoulos C. From Glucose to Lactate and Transiting Intermediates Through Mitochondria, Bypassing Pyruvate Kinase: Considerations for Cells Exhibiting Dimeric PKM2 or Otherwise Inhibited Kinase Activity. Front Physiol 2020;11. DOI:10.3389/fphys.2020.543564.

[56] Zangari J, Petrelli F, Maillot B, Martinou JC. The multifaceted pyruvate metabolism: Role of the mitochondrial pyruvate carrier. Biomolecules 2020;10:1–18. PMID:32708919. DOI:10.3390/biom10071068.

[57] Bittar PG, Charnay Y, Pellerin L, Bouras C, Magistretti PJ. Selective distribution of lactate dehydrogenase isoenzymes in neurons and astrocytes of human brain. J Cereb Blood Flow Metab 1996;16:1079–89. PMID:8898679. DOI:10.1097/00004647-199611000-00001.

[58] Laughton JD, Bittar P, Charnay Y, Pellerin L, Kovari E, Magistretti PJ, et al. Metabolic compartmentalization in the human cortex and hippocampus: evidence for a cell-and region-specific localization of lactate dehydrogenase 5 and pyruvate dehydrogenase. BMC Neurosci 2007;8:35. PMID:17521432. DOI:10.1186/1471-2202-8-35.

[59] Kaplan NO, Ciotti MM, Stolzenbach FE. REACTION OF PYRIDINE NUCLEOTIDE ANALOGUES WITH DEHYDROGENASES. J Biol Chem 1956;221:833–44. PMID:13357478. DOI:10.1016/S0021-9258(18)65197-X.

[60] Quistorff B, Grunnet N. The isoenzyme pattern of LDH does not play a physiological role; except perhaps during fast transitions in energy metabolism. Aging (Albany NY) 2011;3:457–60. PMID:21566263. DOI:10.18632/aging.100329.

[61] Latner AL, Siddiqui SA, Skillen AW. Pyruvate Inhibition of Lactate Dehydrogenase Activity in Human Tissue Extracts. Science (80-) 1966;154:527–9. DOI:10.1126/science.154.3748.527.

[62] Markert CL, Ursprung H. The ontogeny of isozyme patterns of lactate dehydrogenase in the mouse. Dev Biol 1962;5:363–81. DOI:10.1016/0012-1606(62)90019-2.

[63] Vesell ES. Lactate Dehydrogenase Isozymes: Substrate Inhibition in Various Human Tissues. Science (80-) 1965;150:1590–3. DOI:10.1126/science.150.3703.1590.

[64] Pesce A, Fondy TP, Stolzenbach F, Castillo F, Kaplan NO. The Comparative Enzymology of Lactic Dehydrogenases. J Biol Chem 1967;242:2151–67. DOI:10.1016/S0021-9258(18)96030-8.

[65] Kaplan NO, Everse J, Admiraal J. SIGNIFICANCE OF SUBSTRATE INHIBITION OF DEHYDROGENASES. Ann N Y Acad Sci 1968;151:400–12. DOI:10.1111/j.1749-6632.1968.tb11903.x.

[66] Bishop MJ, Everse J, Kaplan NO. Identification of Lactate Dehydrogenase Isoenzymes by Rapid Kinetics. Proc Natl Acad Sci 1972;69:1761–5. PMID:4340158. DOI:10.1073/pnas.69.7.1761.

[67] Kaplan NO, Everse J. Regulatory characteristics of lactate dehydrogenases. Adv Enzyme Regul 1972;10:323–36. PMID:4347315. DOI:10.1016/0065-2571(72)90021-0.

[68] Dawson DM, Goodfriend TL, Kaplan N. Lactic Dehydrogenases: Functions of the Two Types. Science (80-) 1963:929–33. PMID:14090142. DOI:10.1126/science.143.3609.929.

[69] Abu-Shumays RL, Fristrom JW. IMP-L3, a 20-hydroxyecdysone-responsive gene encodes Drosophila lactate dehydrogenase: Structural characterization and developmental studies. Dev Genet 1997;20:11–22. PMID:9094207. DOI:10.1002/(SICI)1520-6408(1997)20:1<11::AID-DVG2>3.0.CO;2-C.

[70] Onoufriou A, Alahiotis SN. Drosophila lactate dehydrogenase: Molecular and genetic aspects. Biochem Genet 1982;20:1195–209. DOI:10.1007/BF00498943.

[71] Charlton-Perkins MA, Sendler ED, Buschbeck EK, Cook TA. Multifunctional glial support by Semper cells in the Drosophila retina. PLOS Genet 2017;13:e1006782. PMID:28562601. DOI:10.1371/journal.pgen.1006782.

[72] Jones SG, Nixon KCJ, Chubak MC, Kramer JM. Mushroom Body Specific Transcriptome Analysis Reveals Dynamic Regulation of Learning and Memory Genes After Acquisition of Long-Term Courtship Memory in Drosophila. G3 Genes|Genomes|Genetics 2018;8:3433–46. PMID:30158319. DOI:10.1534/g3.118.200560.

[73] Brown JB, Boley N, Eisman R, May GE, Stoiber MH, Duff MO, et al. Diversity and dynamics of the Drosophila transcriptome. Nature 2014;512:393–9. PMID:24670639. DOI:10.1038/nature12962.

[74] Scialò F, Sriram A, Fernández-Ayala D, Gubina N, Lõhmus M, Nelson G, et al. Mitochondrial ROS Produced via Reverse Electron Transport Extend Animal Lifespan. Cell Metab 2016;23:725–34. PMID:27076081. DOI:10.1016/j.cmet.2016.03.009.

[75] Hall H, Medina P, Cooper DA, Escobedo SE, Rounds J, Brennan KJ, et al. Transcriptome profiling of aging Drosophila photoreceptors reveals gene expression trends that correlate with visual senescence. BMC Genomics 2017;18:894. PMID:29162050. DOI:10.1186/s12864-017-4304-3.

[76] Hunt LC, Demontis F. Age-Related Increase in Lactate Dehydrogenase Activity in Skeletal Muscle Reduces Lifespan in Drosophila. J Gerontol A Biol Sci Med Sci 2021;XX:1–32. PMID:34477202. DOI:10.1093/gerona/glab260.

[77] Iliadi KG, Boulianne GL. Age-related behavioral changes in Drosophila. Ann N Y Acad Sci 2010;1197:9–18. PMID:20536827. DOI:10.1111/j.1749-6632.2009.05372.x.

[78] Brenman-Suttner DB, Long SQ, Kamesan V, de Belle JN, Yost RT, Kanippayoor RL, et al. Progeny of old parents have increased social space in Drosophila melanogaster. Sci Rep 2018;8:3673. PMID:29487349. DOI:10.1038/s41598-018-21731-0.

[79] Tully T, Quinn WG. Classical conditioning and retention in normal and mutantDrosophila melanogaster. J Comp Physiol A 1985;157:263–77. PMID:3939242. DOI:10.1007/BF01350033.

[80] Mery F. Aging and its differential effects on consolidated memory forms in Drosophila. Exp Gerontol 2007;42:99–101. DOI:10.1016/j.exger.2006.06.004.

[81] Tonoki A, Davis RL. Aging impairs protein-synthesis-dependent long-term memory in Drosophila. J Neurosci 2015;35:1173–80. PMID:25609631. DOI:10.1523/JNEUROSCI.0978-14.2015.

[82] Plaçais P-Y, Preat T. To Favor Survival Under Food Shortage, the Brain Disables Costly Memory. Science (80-) 2013;339:440–2. DOI:10.1126/science.1226018.

[83] Wu C-L, Chang C-C, Wu J-K, Chiang M-H, Yang C-H, Chiang H-C. Mushroom body glycolysis is required for olfactory memory in Drosophila. Neurobiol Learn Mem 2018;150:13–9. PMID:29477608. DOI:10.1016/j.nlm.2018.02.015.

[84] Siegel RW, Hall JC. Conditioned responses in courtship behavior of normal and mutant Drosophila. Proc Natl Acad Sci 1979;76:3430–4. PMID:16592682. DOI:10.1073/pnas.76.7.3430.

[85] McBride SM., Giuliani G, Choi C, Krause P, Correale D, Watson K, et al. Mushroom Body Ablation Impairs Short-Term Memory and Long-Term Memory of Courtship Conditioning in Drosophila melanogaster. Neuron 1999;24:967–77. PMID:10624959. DOI:10.1016/S0896-6273(00)81043-0.

[86] Heisenberg M, Borst A, Wagner S, Byers D. Drosophila Mushroom Body Mutants are Deficient in Olfactory Learning. J Neurogenet 1985;2:1–30. PMID:4020527. DOI:10.3109/01677068509100140.

[87] Tully T, Preat T, Boynton SC, Del Vecchio M. Genetic dissection of consolidated memory in Drosophila. Cell 1994;79:35–47. PMID:7923375. DOI:10.1016/0092-8674(94)90398-0.

[88] Savvateeva E V, Popov A V, Kamyshev NG, Iliadi KG, Bragina J V, Heisenberg M, et al. Age-dependent changes in memory and mushroom bodies in the Drosophila mutant vermilion deficient in the kynurenine pathway of tryptophan metabolism. Russ J Physiol 1999;85:167–83. PMID:10389174..

[89] McGuire SE, Le PT, Osborn AJ, Matsumoto K, Davis RL. Spatiotemporal Rescue of Memory Dysfunction in Drosophila. Science (80-) 2003;302:1765–8. PMID:14657498. DOI:10.1126/science.1089035.

[90] Brown EJ, Nguyen AH, Bachtrog D. The Y chromosome may contribute to sex-specific ageing in Drosophila. Nat Ecol Evol 2020;4:853–62. PMID:32313175. DOI:10.1038/s41559-020-1179-5.

[91] Smolič T, Tavčar P, Horvat A, Černe U, Halužan A, Tratnjek L, et al. Astrocytes in stress accumulate lipid droplets. Glia 2021;69:1540–62. DOI:10.1002/glia.23978.

[92] Greenspan P, Mayer EP, Fowler SD. Nile red: A selective fluorescent stain for intracellular lipid droplets. J Cell Biol 1985;100:965–73. PMID:3972906. DOI:10.1083/jcb.100.3.965.

[93] Chen M, Sokolowski MB. How Social Experience and Environment Impacts Behavioural Plasticity in Drosophila. Fly (Austin) 2022;16:68–84. PMID:34852730. DOI:10.1080/19336934.2021.1989248.

[94] Leech T, Sait SM, Bretman A. Sex-specific effects of social isolation on ageing in Drosophila melanogaster. J Insect Physiol 2017;102:12–7. PMID:28830760. DOI:10.1016/j.jinsphys.2017.08.008.

[95] Vora A, Nguyen AD, Spicer C, Li W. The impact of social isolation on health and behavior in Drosophila melanogaster and beyond. Brain Sci Adv 2022;8:183–96. DOI:10.26599/BSA.2022.9050016.

[96] Sunderhaus ER, Kretzschmar D. Mass Histology to Quantify Neurodegeneration in <em>Drosophila</em>. J Vis Exp 2016;2016:1–7. PMID:28060320. DOI:10.3791/54809.

[97] Park JS, Saeed K, Jo MH, Kim MW, Lee HJ, Park C-B, et al. LDHB Deficiency Promotes Mitochondrial Dysfunction Mediated Oxidative Stress and Neurodegeneration in Adult Mouse Brain. Antioxidants 2022;11:261. DOI:10.3390/antiox11020261.

[98] Phillips MA, Arnold KR, Vue Z, Beasley HK, Garza-Lopez E, Marshall AG, et al. Combining Metabolomics and Experimental Evolution Reveals Key Mechanisms Underlying Longevity Differences in Laboratory Evolved Drosophila melanogaster Populations. Int J Mol Sci 2022;23:1067. PMID:35162994. DOI:10.3390/ijms23031067.

[99] Westfall S, Lomis N, Prakash S. Longevity extension in Drosophila through gut-brain communication. Sci Rep 2018;8:1–15. PMID:29849035. DOI:10.1038/s41598-018-25382-z.

[100] López-Otín C, Blasco MA, Partridge L, Serrano M, Kroemer G. The Hallmarks of Aging. Cell 2013;153:1194–217. PMID:23746838. DOI:10.1016/j.cell.2013.05.039.

[101] Sonn JY, Lee J, Sung MK, Ri H, Choi JK, Lim C, et al. Serine metabolism in the brain regulates starvation-induced sleep suppression in Drosophila melanogaster. Proc Natl Acad Sci 2018;115:7129–34. PMID:29915051. DOI:10.1073/pnas.1719033115.

[102] Dai X, Zhou E, Yang W, Zhang X, Zhang W, Rao Y. D-Serine made by serine racemase in Drosophila intestine plays a physiological role in sleep. Nat Commun 2019;10:1986. PMID:31064979. DOI:10.1038/s41467-019-09544-9.

[103] Suzuki M, Sasabe J, Miyoshi Y, Kuwasako K, Muto Y, Hamase K, et al. Glycolytic flux controls <scp>d</scp>-serine synthesis through glyceraldehyde-3-phosphate dehydrogenase in astrocytes. Proc Natl Acad Sci 2015;112:E2217–24. PMID:25870284. DOI:10.1073/pnas.1416117112.

[104] Le Douce J, Maugard M, Veran J, Matos M, Jégo P, Vigneron P-A, et al. Impairment of Glycolysis-Derived l-Serine Production in Astrocytes Contributes to Cognitive Deficits in Alzheimer’s Disease. Cell Metab 2020;31:503–517.e8. PMID:32130882. DOI:10.1016/j.cmet.2020.02.004.

[105] Kurniawan H, Franchina DG, Guerra L, Bonetti L, Baguet LS, Grusdat M, et al. Glutathione Restricts Serine Metabolism to Preserve Regulatory T Cell Function. Cell Metab 2020;31:920–936.e7. PMID:32213345. DOI:10.1016/j.cmet.2020.03.004.

[106] Newman AC, Maddocks ODK. One-carbon metabolism in cancer. Br J Cancer 2017;116:1499–504. PMID:28472819. DOI:10.1038/bjc.2017.118.

[107] Ducker GS, Rabinowitz JD. One-Carbon Metabolism in Health and Disease. Cell Metab 2017;25:27–42. PMID:27641100. DOI:10.1016/j.cmet.2016.08.009.

[108] Coyle JT, Balu D, Wolosker H. d-Serine, the Shape-Shifting NMDA Receptor Co-agonist. Neurochem Res 2020;45:1344–53. PMID:32189130. DOI:10.1007/s11064-020-03014-1.

[109] Graham C, Stefanatos R, Yek AEH, Spriggs R V, Loh SHY, Uribe AH, et al. Mitochondrial ROS signalling requires uninterrupted electron flow and is lost during ageing in flies. GeroScience 2022. DOI:10.1007/s11357-022-00555-x.

[110] Sanz A, Stefanatos R, McIlroy G. Production of reactive oxygen species by the mitochondrial electron transport chain in Drosophila melanogaster. J Bioenerg Biomembr 2010;42:135–42. PMID:20300811. DOI:10.1007/s10863-010-9281-z.

[111] Linnane A, Ozawa T, Marzuki S, Tanaka M. MITOCHONDRIAL DNA MUTATIONS AS AN IMPORTANT CONTRIBUTOR TO AGEING AND DEGENERATIVE DISEASES. Lancet 1989;333:642–5. PMID:2564461. DOI:10.1016/S0140-6736(89)92145-4.

[112] Tatar M. Can We Develop Genetically Tractable Models to Assess Healthspan (Rather Than Life Span) in Animal Models? Journals Gerontol Ser A Biol Sci Med Sci 2009;64A:161–3. PMID:19225031. DOI:10.1093/gerona/gln067.

[113] Mahmoudzadeh NH, Fitt AJ, Schwab DB, Martenis WE, Nease LM, Owings CG, et al. The oncometabolite L-2-hydroxyglutarate is a common product of dipteran larval development. Insect Biochem Mol Biol 2020;127:103493. PMID:33157229. DOI:10.1016/j.ibmb.2020.103493.

[114] Li H, Chawla G, Hurlburt AJ, Sterrett MC, Zaslaver O, Cox J, et al. Drosophila larvae synthesize the putative oncometabolite L-2-hydroxyglutarate during normal developmental growth. Proc Natl Acad Sci 2017;114:1353–8. PMID:28115720. DOI:10.1073/pnas.1614102114.

[115] Zhao X, Golic FT, Harrison BR, Manoj M, Hoffman E V., Simon N, et al. The metabolome as a biomarker of aging in Drosophila melanogaster. Aging Cell 2022;21:1–14. PMID:35019203. DOI:10.1111/acel.13548.

[116] Hunt RJ, Granat L, McElroy GS, Ranganathan R, Chandel NS, Bateman JM. Mitochondrial stress causes neuronal dysfunction via an ATF4-dependent increase in L-2-hydroxyglutarate. J Cell Biol 2019;218:4007–16. PMID:31645461. DOI:10.1083/jcb.201904148.

[117] Fu X, Chin RM, Vergnes L, Hwang H, Deng G, Xing Y, et al. 2-Hydroxyglutarate Inhibits ATP Synthase and mTOR Signaling. Cell Metab 2015;22:508–15. PMID:26190651. DOI:10.1016/j.cmet.2015.06.009.

[118] Brinkley G, Nam H, Shim E, Kirkman R, Kundu A, Karki S, et al. Teleological Role of L-2-Hydroxyglutarate Dehydrogenase in the Kidney. Dis Model Mech 2020;13. PMID:32928875. DOI:10.1242/dmm.045898.

[119] Rodrigues DGB, de Moura Coelho D, Sitta Â, Jacques CED, Hauschild T, Manfredini V, et al. Experimental evidence of oxidative stress in patients with l-2-hydroxyglutaric aciduria and that l-carnitine attenuates in vitro DNA damage caused by d-2-hydroxyglutaric and l-2-hydroxyglutaric acids. Toxicol Vitr 2017;42:47–53. PMID:28396261. DOI:10.1016/j.tiv.2017.04.006.

[120] Kranendijk M, Struys EA, Salomons GS, Van der Knaap MS, Jakobs C. Progress in understanding 2-hydroxyglutaric acidurias. J Inherit Metab Dis 2012;35:571–87. PMID:22391998. DOI:10.1007/s10545-012-9462-5.

[121] Ma S, Sun R, Jiang B, Gao J, Deng W, Liu P, et al. L2hgdh Deficiency Accumulates l-2-Hydroxyglutarate with Progressive Leukoencephalopathy and Neurodegeneration. Mol Cell Biol 2017;37:550. DOI:10.1128/MCB.00492-16.

[122] Johnson AA, Stolzing A. The role of lipid metabolism in aging, lifespan regulation, and age-related disease. Aging Cell 2019;18:1–26. PMID:31560163. DOI:10.1111/acel.13048.

[123] Welte MA, Gould AP. Lipid droplet functions beyond energy storage. Biochim Biophys Acta-Mol Cell Biol Lipids 2017;1862:1260–72. PMID:28735096. DOI:10.1016/j.bbalip.2017.07.006.

[124] Cohen S. Lipid Droplets as Organelles. Int. Rev. Cell Mol. Biol., vol. 337. 1st ed., Elsevier Inc.; 2018, p. 83–110. PMID:29551163. DOI:10.1016/bs.ircmb.2017.12.007.

[125] Ralhan I, Chang CL, Lippincott-Schwartz J, Ioannou MS. Lipid droplets in the nervous system. J Cell Biol 2021;220:1–18. PMID:34152362. DOI:10.1083/jcb.202102136.

[126] Zechner R, Zimmermann R, Eichmann TO, Kohlwein SD, Haemmerle G, Lass A, et al. FAT SIGNALS-Lipases and Lipolysis in Lipid Metabolism and Signaling. Cell Metab 2012;15:279–91. PMID:22405066. DOI:10.1016/j.cmet.2011.12.018.

[127] Kis V, Barti B, Lippai M, Sass M. Specialized Cortex Glial Cells Accumulate Lipid Droplets in Drosophila melanogaster. PLoS One 2015;10:e0131250. PMID:26148013. DOI:10.1371/journal.pone.0131250.

[128] Dong Q, Zavortink M, Froldi F, Golenkina S, Lam T, Cheng LY. Glial Hedgehog signalling and lipid metabolism regulate neural stem cell proliferation in Drosophila. EMBO Rep 2021;22:1–18. DOI:10.15252/embr.202052130.

[129] Stenger C, Pinçon A, Hanse M, Royer L, Comte A, Koziel V, et al. Brain region-specific immunolocalization of the lipolysis-stimulated lipoprotein receptor (LSR) and altered cholesterol distribution in aged LSR +/-mice. J Neurochem 2012;123:467–76. PMID:22909011. DOI:10.1111/j.1471-4159.2012.07922.x.

[130] Shimabukuro MK, Langhi LGP, Cordeiro I, Brito JM, Batista CMDC, Mattson MP, et al. Lipid-laden cells differentially distributed in the aging brain are functionally active and correspond to distinct phenotypes. Sci Rep 2016;6:1–12. PMID:27029648. DOI:10.1038/srep23795.

[131] Marschallinger J, Iram T, Zardeneta M, Lee SE, Lehallier B, Haney MS, et al. Lipid-droplet-accumulating microglia represent a dysfunctional and proinflammatory state in the aging brain. Nat Neurosci 2020;23:194–208. PMID:31959936. DOI:10.1038/s41593-019-0566-1.

[132] Liu L, Zhang K, Sandoval H, Yamamoto S, Jaiswal M, Sanz E, et al. Glial lipid droplets and ROS induced by mitochondrial defects promote neurodegeneration. Cell 2015;160:177–90. PMID:25594180. DOI:10.1016/j.cell.2014.12.019.

[133] Bailey AP, Koster G, Guillermier C, Hirst EMA, MacRae JI, Lechene CP, et al. Antioxidant Role for Lipid Droplets in a Stem Cell Niche of Drosophila. Cell 2015;163:340–53. PMID:26451484. DOI:10.1016/j.cell.2015.09.020.

[134] Girard V, Goubard V, Querenet M, Seugnet L, Pays L, Nataf S, et al. Spen modulates lipid droplet content in adult Drosophila glial cells and protects against paraquat toxicity. Sci Rep 2020;10:20023. PMID:33208773. DOI:10.1038/s41598-020-76891-9.

[135] Fei W, Wang H, Fu X, Bielby C, Yang H. Conditions of endoplasmic reticulum stress stimulate lipid droplet formation in Saccharomyces cerevisiae. Biochem J 2009;424:61–7. PMID:19708857. DOI:10.1042/BJ20090785.

[136] Velázquez AP, Tatsuta T, Ghillebert R, Drescher I, Graef M. Lipid droplet–mediated ER homeostasis regulates autophagy and cell survival during starvation. J Cell Biol 2016;212:621–31. PMID:26953354. DOI:10.1083/jcb.201508102.

[137] Farmer BC, Walsh AE, Kluemper JC, Johnson LA. Lipid Droplets in Neurodegenerative Disorders. Front Neurosci 2020;14:1–13. DOI:10.3389/fnins.2020.00742.

[138] Goodman LD, Bellen HJ. Recent insights into the role of glia and oxidative stress in Alzheimer’s disease gained from Drosophila. Curr Opin Neurobiol 2022;72:32–8. PMID:34418791. DOI:10.1016/j.conb.2021.07.012.

[139] Moulton MJ, Barish S, Ralhan I, Chang J, Goodman LD, Harland JG, et al. Neuronal ROS-induced glial lipid droplet formation is altered by loss of Alzheimer’s disease–associated genes. Proc Natl Acad Sci 2021;118:e2112095118. PMID:34949639. DOI:10.1073/pnas.2112095118.

[140] Alzheimer A. über eigenartige Krankheitsfälle des späteren Alters. Zeitschrift Für Die Gesamte Neurol Und Psychiatr 1911;4:356–85. DOI:10.1007/BF02866241.

[141] Wat LW, Chao C, Bartlett R, Buchanan JL, Millington JW, Chih HJ, et al. A role for triglyceride lipase brummer in the regulation of sex differences in Drosophila fat storage and breakdown. PLOS Biol 2020;18:e3000595. PMID:31961851. DOI:10.1371/journal.pbio.3000595.

[142] Yang C, Wang X, Wang J, Wang X, Chen W, Lu N, et al. Rewiring Neuronal Glycerolipid Metabolism Determines the Extent of Axon Regeneration. Neuron 2020;105:276–292.e5. PMID:31786011. DOI:10.1016/j.neuron.2019.10.009.

[143] Schönfeld P, Reiser G. Brain energy metabolism spurns fatty acids as fuel due to their inherent mitotoxicity and potential capacity to unleash neurodegeneration. Neurochem Int 2017;109:68–77. PMID:28366720. DOI:10.1016/j.neuint.2017.03.018.

[144] Schönfeld P, Reiser G. Why does Brain Metabolism not Favor Burning of Fatty Acids to Provide Energy?-Reflections on Disadvantages of the Use of Free Fatty Acids as Fuel for Brain. J Cereb Blood Flow Metab 2013;33:1493–9. PMID:23921897. DOI:10.1038/jcbfm.2013.128.

[145] Yin J, Spillman E, Cheng ES, Short J, Chen Y, Lei J, et al. Brain-specific lipoprotein receptors interact with astrocyte derived apolipoprotein and mediate neuron-glia lipid shuttling. Nat Commun 2021;12:2408. DOI:10.1038/s41467-021-22751-7.

[146] Ioannou MS, Jackson J, Sheu SH, Chang CL, Weigel A V., Liu H, et al. Neuron-Astrocyte Metabolic Coupling Protects against Activity-Induced Fatty Acid Toxicity. Cell 2019;177:1522–1535.e14. PMID:31130380. DOI:10.1016/j.cell.2019.04.001.

[147] Qi G, Mi Y, Shi X, Gu H, Brinton RD, Yin F. ApoE4 Impairs Neuron-Astrocyte Coupling of Fatty Acid Metabolism. Cell Rep 2021;34:108572. PMID:33406436. DOI:10.1016/j.celrep.2020.108572.

[148] Harris RA, Lone A, Lim H, Martinez F, Frame AK, Scholl TJ, et al. Aerobic Glycolysis Is Required for Spatial Memory Acquisition But Not Memory Retrieval in Mice. Eneuro 2019;6:ENEURO.0389-18.2019. DOI:10.1523/ENEURO.0389-18.2019.

[149] Jensen CJ, Demol F, Bauwens R, Kooijman R, Massie A, Villers A, et al. Astrocytic β2 Adrenergic Receptor Gene Deletion Affects Memory in Aged Mice. PLoS One 2016;11:e0164721. DOI:10.1371/journal.pone.0164721.

[150] Raun N, Jones S, Kramer JM. Conditioned courtship suppression in Drosophila melanogaster. J Neurogenet 2021;0:1–27. DOI:10.1080/01677063.2021.1873323.

[151] Zyla RE, Hodgson A. Gene of the month: FH. J Clin Pathol 2021;74:615–9. PMID:34353877. DOI:10.1136/jclinpath-2021-207830.

[152] Coman D, Kranc KR, Christodoulou J. Fumarate Hydratase Deficiency. 1993. PMID:20301679..

[153] Koemans TS, Kleefstra T, Chubak MC, Stone MH, Reijnders MRF, de Munnik S, et al. Functional convergence of histone methyltransferases EHMT1 and KMT2C involved in intellectual disability and autism spectrum disorder. PLOS Genet 2017;13:e1006864. DOI:10.1371/journal.pgen.1006864.

[154] Garner SRC, Castellanos MC, Baillie K, Lian T, Allan DW. Drosophila female-specific Ilp7 motoneurons are generated by Fruitless-dependent cell death in males and a double-assurance survival role for Transformer in females. Development 2017;145. PMID:29229771. DOI:10.1242/dev.150821.

[155] Schindelin J, Arganda-Carreras I, Frise E, Kaynig V, Longair M, Pietzsch T, et al. Fiji: an open-source platform for biological-image analysis. Nat Methods 2012;9:676–82. DOI:10.1038/nmeth.2019.

[156] Piper MDW, Partridge L. Protocols to Study Aging in Drosophila. Methods Mol. Biol., vol. 1478, 2016, p. 291–302. PMID:27730590. DOI:10.1007/978-1-4939-6371-3_18.

[157] Li H, Tennessen JM. Preparation of <em>Drosophila</em> Larval Samples for Gas Chromatography-Mass Spectrometry (GC-MS)-based Metabolomics. J Vis Exp 2018;2018:1–7. PMID:29939167. DOI:10.3791/57847.

[158] Motulsky HJ, Brown RE. Detecting outliers when fitting data with nonlinear regression-a new method based on robust nonlinear regression and the false discovery rate. BMC Bioinformatics 2006;7:123. PMID:16526949. DOI:10.1186/1471-2105-7-123.

[159] McNeil AR, Jolley SN, Akinleye AA, Nurilov M, Rouzyi Z, Milunovich AJ, et al. Conditions Affecting Social Space in <em>Drosophila melanogaster</em>. J Vis Exp 2015;2015:1–10. PMID:26575105. DOI:10.3791/53242.

[160] Benzer S. BEHAVIORAL MUTANTS OF Drosophila ISOLATED BY COUNTERCURRENT DISTRIBUTION. Proc Natl Acad Sci 1967;58:1112–9. PMID:16578662. DOI:10.1073/pnas.58.3.1112.

[161] Shihan MH, Novo SG, Le Marchand SJ, Wang Y, Duncan MK. A simple method for quantitating confocal fluorescent images. Biochem Biophys Reports 2021;25:100916. DOI:10.1016/j.bbrep.2021.100916.

